# Mitochondria form contact sites with the nucleus to couple pro-survival retrograde response

**DOI:** 10.1101/445411

**Authors:** Radha Desai, Daniel A East, Liana Hardy, James Crosby, Manuel Rigon, Danilo Faccenda, María Soledad Alvarez, Aarti Singh, Marta Mainenti, Laura Kuhlman Hussey, Robert Bentham, Gyorgy Szabadkai, Valentina Zappulli, Gurtej Dhoot, Lisa E Romano, Xia Dong, Isabelle Coppens, Anne Hamacher-Brady, J Paul Chapple, Rosella Abeti, Roland A. Fleck, Gema Vizcay-Barrena, Kenneth Smith, Michelangelo Campanella

## Abstract

Mitochondria drive cellular adaptation to stress by retro-communicating with the nucleus. This process is known as Mitochondrial Retrograde Response (MRR) and is induced by mitochondrial dysfunctions which perturb cell signalling. MRR results in the nuclear stabilization and activation of pro-survival transcription factors such as the nuclear factor kappa-light-chain-enhancer of activated B cells (NF-kB). Here we demonstrate that MRR is facilitated by the formation of contact sites between mitochondria and the nucleus which establish microdomains of communication between the two organelles. The 18kD Translocator Protein (TSPO), which de-ubiquitylates and stabilizes the mitochondrial network preventing its mitophagy-mediated segregation, is required for this interaction. The tethering TSPO enacts is mediated by the complex formed with the Protein Kinase A via the A-kinase anchoring protein Acyl-CoA Binding Domain Containing 3 (ACBD3) and allows the redistribution of cholesterol which sustains the pro-survival response by blocking NF-kB de-acetylation. This work proposes a new paradigm in the mitochondrial retro-communication by revealing the existence of contact sites between mitochondrial and the nucleus and a signalling role for cholesterol.

## Introduction

Mitochondria actively participate in the reprogramming of mammalian cells^1,2^. In response to endogenous or exogenous perturbations they retro-communicate with the nucleus to drive the transcription of genes. This pathway of signalling is known as the Mitochondrial Retrograde Response (MRR) and is exploited in the pathogenesis of uncontrolled cellular proliferation^3^. In cancer pro-survival transcription factors (TFs) like the nuclear factor kappa-light-chain-enhancer of activated B cells (NF-kB^4–7^) are stabilised by the hyper-production of Reactive Oxygen Species (ROS) generated from dysfunctional mitochondria. Impaired cell signalling and nuclear accumulation of NF-kB are the acknowledged read-outs of active MRR in mammals. In these species, the lack of homology with the yeast Rtg1/Rtg2/Rtg3 system^10^, entails the engagement of a more complex framework of regulators and executors^11^ when mitochondria communicate with the nucleus. In keeping with this, the recent identification of MRR molecules^8,9^ failed to explain the interplay between mitochondria and nucleus during stress. Aspects intimately involved in the organelle physiopathology such as dynamic remodelling (i) and trafficking of lipids (ii) had been unexplored and for these we postulated a catalytic role in the MRR signalling. Abnormalities in lipid metabolism are distinctive features of cancer cells which vastly rely on MRR^12^ and in these the increased mitochondrial network size following the stabilization of pro-fusion proteins (e.g. OPA-1, Mnf1, 2, ATPIF1^13–15^) mediates cytoprotective resistance to organelle disassembly. Hypertrophic mitochondria by occupying the surrounding cytosolic space would therefore hold greater chance of establishing proximity events with other organelles thus generating key hotspots of communication. It is indeed well established that hyperfused, remodelled mitochondria interact with the Endoplasmic Reticulum (ER) whilst unexplored is whether they become physically coupled with the nucleus. If this was proved correct it would be represent a prerequisite for the activation and execution of MRR.

In order to test the hypothesis of domains of communication between mitochondria and nucleus during MRR which redistribute cholesterol, which is eponymous of the lipid dysregulation, we traced the molecular physiology of the Outer Mitochondrial Membrane (OMM) based protein TSPO^16^. This stress-responsive protein is upregulated in many cancers^17,18^ and linked to anti-apoptotic activity. Its pharmacological ligands allow its selective targeting to co-adjuvate chemotherapy and *in vivo* tracing of metastatic cells ^20,21^. Once accumulated on the OMM, TSPO represses mitophagy^22^ and deregulates mitochondrial buffering capacity for Ca^2+^ and production of ATP^23^ originated from the oxidative phosphorylation. TSPO recruits and docks on mitochondria the A-kinase anchoring protein Acyl-CoA Binding Domain Containing 3 (ACBD3) and the holoenzyme Protein kinase A (PKA) 23,24 which complexes with the nucleus via the A-kinase-anchoring protein AKAP95^25,26^. Interestingly, breast cancer in which TSPO is overexpressed^27^ ^28^ have modified signalling^29^ and metabolism^30^ of cholesterol which is trafficked by TSPO to fuel the synthesis of pregnenolone^31–32^ within the mitochondrial matrix^33^. Cholesterol, its intermediates and their enzymatic catalysts such as the aromatase enzyme CYP19A1 are involved in the oncogenic reprogramming^34^ ^35^ of the mammary gland cells. The all which makes TSPO an ideal means to insight the interplay between organelle morphology and lipid metabolism in the mitochondrial retrograde response.

Using cellular and ultrastructural means of analysis combined with advanced protocols of imaging, we show that TSPO is required to establish contact sites between mitochondria and nucleus during MRR allowing for redistribution of cholesterol which de-acetylates NF-kB completing the pro-survival response. A novel regulatory axis in the mitochondrial retrograde response which operates via the formation of contact sites (i) and redistribution of cholesterol (ii) between mitochondria and nucleus is therefore outlined. These evidences advance our understanding of a key process in mammalian pathophysiology and detail a physical and metabolic coupling between mitochondria and nucleus previously unknown.

## Results

### Mitochondrial retrograde response associates with organelle remodelling and high of level of TSPO expression which promotes NF-kB stabilization into the nucleus

TSPO expression, which positively associates with breast cancer aggressiveness, drives NF-κB nuclear recruitment. The first step of our analysis consisted in assessing the correlation between the candidate protein TSPO and the nuclear translocation of the pro-survival transcriptional factor NF-κB^36,37^, which is an acknowledged read-out of MRR. We therefore inspected primary human samples of mammary gland tumours reporting a positive association between TSPO level, aggressiveness of the lesions (**Figure 1 a, b**), and NF-κB accumulation into the nucleus (**Figure 1d, e**). In line with this, transcriptomic analysis of >600 samples of breast cancer patients in the Cancer Genome Atlas (TCGA) showed higher levels of active NF-κB correlated with increased TSPO expression in the more aggressive forms of tumours (HER positive and basal types) (**Figure 1c, f**), without detecting any mutation in TSPO (**SFigure 1a**). The increased expression of TSPO as underlying feature in the neoplasia of the mammary gland is notably retained across species (**SFigure 1d-g**).

**Figure 1.**
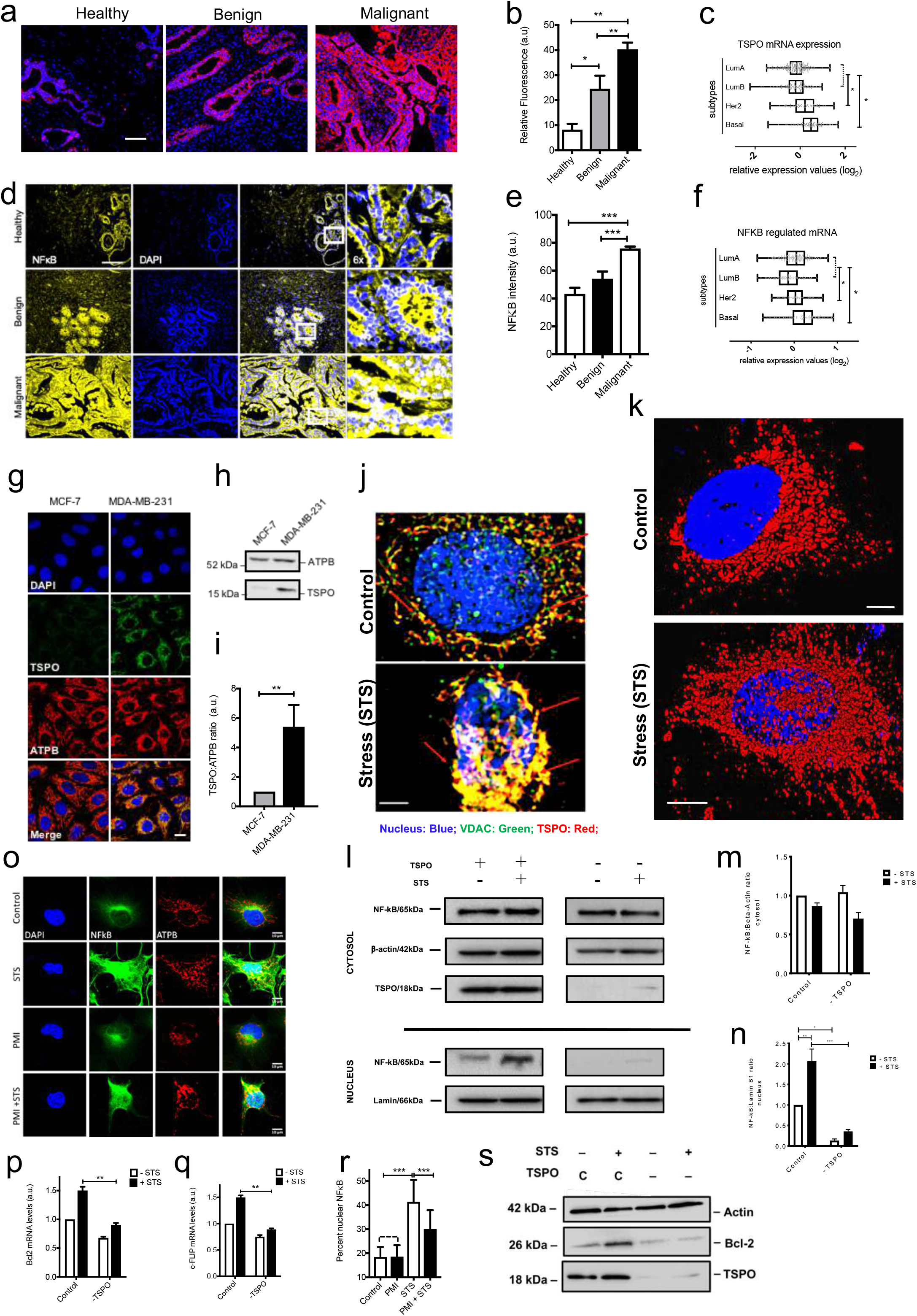
TSPO rich mitochondria gather on the nucleus during apoptotic treatment and dictate the degree of NF-kB retro-translocation. (a) Pseudo-coloured images of fluorescent immunohistochemistry using anti-TSPO antibody (red) reveals higher expression in human mammary cancer tissue compared with healthy. Nuclei were stained with DAPI. (b) Quantification of TSPO fluorescence normalised to DAPI staining shows difference between cancerous and healthy tissues is statistically significant (Healthy 8.18±2.4; Benign 24.54±5.2; Malignant 40.31±2.7; n=10; p<0.05, 0.01). (c) Comparison of TSPO expression in human mammary tumours using microarray data from the TCGA breast cancer database. (d) Fluorescence immunohistochemical analysis of nuclear NF-κB levels in tissue sections of human mammary gland at various degrees of breast cancer. Samples were immunolabelled for NF-κB (yellow) and nuclei were stained with DAPI (blue). Images show increased expression and translocation of NF-kB into nucleus in malignant samples. (e) The intensity of NF-κB fluorescence shows significantly increased labelling in malignant tissues when compared with benign and healthy tissues (Healthy 43.288±10.6; Benign 54.17±12.7; Malignant 75.94±10.7; n=60 p<0.001) (f) Comparison of NF-κB expression in human mammary tumours using microarray data from the TCGA breast cancer database. (g) Immunohistochemistry of MCF-7 and MDA cells stained for TSPO (green) and ATPB (red) showing higher expression of TSPO in MDA cells. Nuclei were stained with DAPI (blue). (h) Western blot analysis of TSPO levels in MBA and MCF-7 cells (N=4). (i) Quantification of TSPO band density (normalized to ATPB) in western blots shows higher expression of the protein in MDA cells when compared to MCF-7 cells (fold-increase in MDA cells: 5.404004 ± 1.491815, p<0.05) n=3). (j) Confocal microscopy analysis of mitochondrial network remodelling in MDA cells exposed to STS. Immunostaining was performed using anti-VDAC (green) and anti-TSPO (red) antibodies. (k) 3D reconstruction from z-stack confocal images of mitochondrial network remodelling in MDA cells exposed to STS. Mitochondria were stained with anti-TSPO (red) antibody. (l) Western blot analysis of NF-κB levels in the cytosolic and nuclear fractions of MDA cells following STS treatment. Nuclear translocation is significantly reduced in - TSPO cells. (m) Quantification of cytosolic NF-κB normalised to β-actin in control and -TSPO MDA cells treated with STS (Control - STS normalised 1; Control + STS 0.935±0.793; - TSPO - STS 1.193±0.895; - TSPO + STS 0.841±0.567; n=3 p>0.05). (n) Quantification of nuclear NF-κB normalised to lamin B1 in control and -TSPO cells treated with STS (Control - STS normalised 1; Control + STS 2.577±1.557; - TSPO - STS 0.193±0.062; - TSPO + STS 0.435±0.277; n=3 p<0.05, p<0.01, p<0.001). (o) Reduced Bcl-2 mRNA expression in -TSPO MDA cells (Control normalised 1; Control + STS 1.50±0.065; - TSPO 0.68±0.02; - TSPO + STS 0.9±0.04; n=3; p<0.01) (p) Reduced c-FLIP mRNA expression in - TSPO MDA cells (Control normalised 1; Control + STS 1.5±0.04; - TSPO 0.75±0.03; - TSPO + STS 0.89±0.02; n=3; p<0.01). (q) Western blot analysis of Bcl2 levels in control and - TSPO MDA cells following STS treatment. Bcl2 upregulation is not observed in STS-treated, TSPO knock-down cells. (r) Confocal images of fluorescence immunohistochemistry in MDA cells labelled for NF-κB (green) and ATPB (red), treated with STS, PMI and STS+PMI. Nuclei were stained with DAPI (blue). (s) Bar chart of percentage of NF-κB localizing to the nucleus in MDA cells, showing significant nuclear translocation with STS treatment, which is reduced when PMI is introduced (Control 18.42±4.17; PMI 41.39±9.09; STS 18.57±4.7; PMI + STS 30.08±7.8; n=30; p<0.001).

In order to elucidate the nature of this oncogenic interplay between TSPO and NF-κB we moved the analysis *in vitro* and compared epithelial human breast cancer MCF-7 cells, which feature low TSPO level, and the more aggressive counterpart MDA-MB-231 (henceforth referred to as MDA) which instead express high TSPO^29^ level (**Figure 1g-h**). In MDA cells, TSPO was transiently knocked down (-TSPO) (**SFigure 1b**) before exposure to Staurosporine (STS), which was used as a mitochondrial stressor to engage the mitochondrial retrograde response. STS remodels mitochondria on the nucleus (**Figure 1j, k**) and by stabilising the expression of TSPO (**SFigure 1c**) facilitates the nuclear accumulation of NF-κB (**Figure 1l-n**). The outcome of the increased NF-κB retro-translocation was corroborated by analyses of the transcriptional profile of the NF-κB-regulated pro-survival genes c-FLIP and Bcl-2 (**Figure 1o, p**) and protein expression of the latter (**Figure 1q**). Immunocytochemical analysis further highlighted the degree of nuclear residency of NF-κB upon STS treatment and how pruning of the mitochondrial network via pharmacological activation of mitophagy by the p62/SQSTM1-mediated mitophagy inducer (PMI)^38,39^ reduces the degree of NF-κB translocation (**Figure 1r, s**). By reducing the size of the mitochondrial network, PMI limits the cytosolic space occupancy by mitochondria (**SFigure 1j**) and sensitizes human, feline and canine breast cancer cells to STS-induced apoptosis (**SFigure 1k-m**). The integrity of the mitochondrial network, which is reflected by TSPO overexpression and accumulation on the OMM, is therefore pivotal to stabilise the retro-translocation and transcriptomic capacity of NF-κB during MRR activation.

### Targeting of TSPO and its cholesterol binding capacity dictates susceptibility of cells to programmed cell death

Treatment of MDA cells with the TSPO ligand PK11195 (**Table 1**), which competes with the cholesterol binding domains of the protein^40^, inhibits NF-κB retro-translocation similarly to PMI (**Figure 2a, b**). Notably, the cholesterol synthesis inhibitor pravastatin (PVS) counteracts STS mediated effect **(Figure 2c).** This implies the existence of a regulatory hierarchy between TSPO, cholesterol and NF-κB in regulating MRR-mediated evasion of cell death. Confirmation of this was provided by enrolling MDA cells stably knocked-out for NF-κB (NF-kB KO)^41^, in which the additive affect exercised by the concomitant repression of TSPO was deemed (**Figure 2d**). Knocking down of TSPO (-TSPO) in MDA cells competent for NF-kB was indeed able to: increase sensitivity to apoptosis (i) (**Figure 2e, SFigure 1h,i**), reduce cellular proliferative capacity (ii) (**Figure 2f**), and facilitate the accumulation of the pro-apoptotic protein Bax on mitochondria (iii) (**Figure 2g,h**), as well as release of cytochrome C in the cytosol (iv) (**Figure 2i,j**). In keeping with this, TSPO ligands (**Table 1**) tangibly promoted susceptibility to chemically induced cell death, thereby co-adjuvating not only the effect of STS treatment (**Figure 2k**), but also that of Doxorubicin or Vincristine (**Figure 2l,m**) in both two-dimensional and in three-dimensional cell cultures (**SFigure 3c)**.

**Figure 2.**
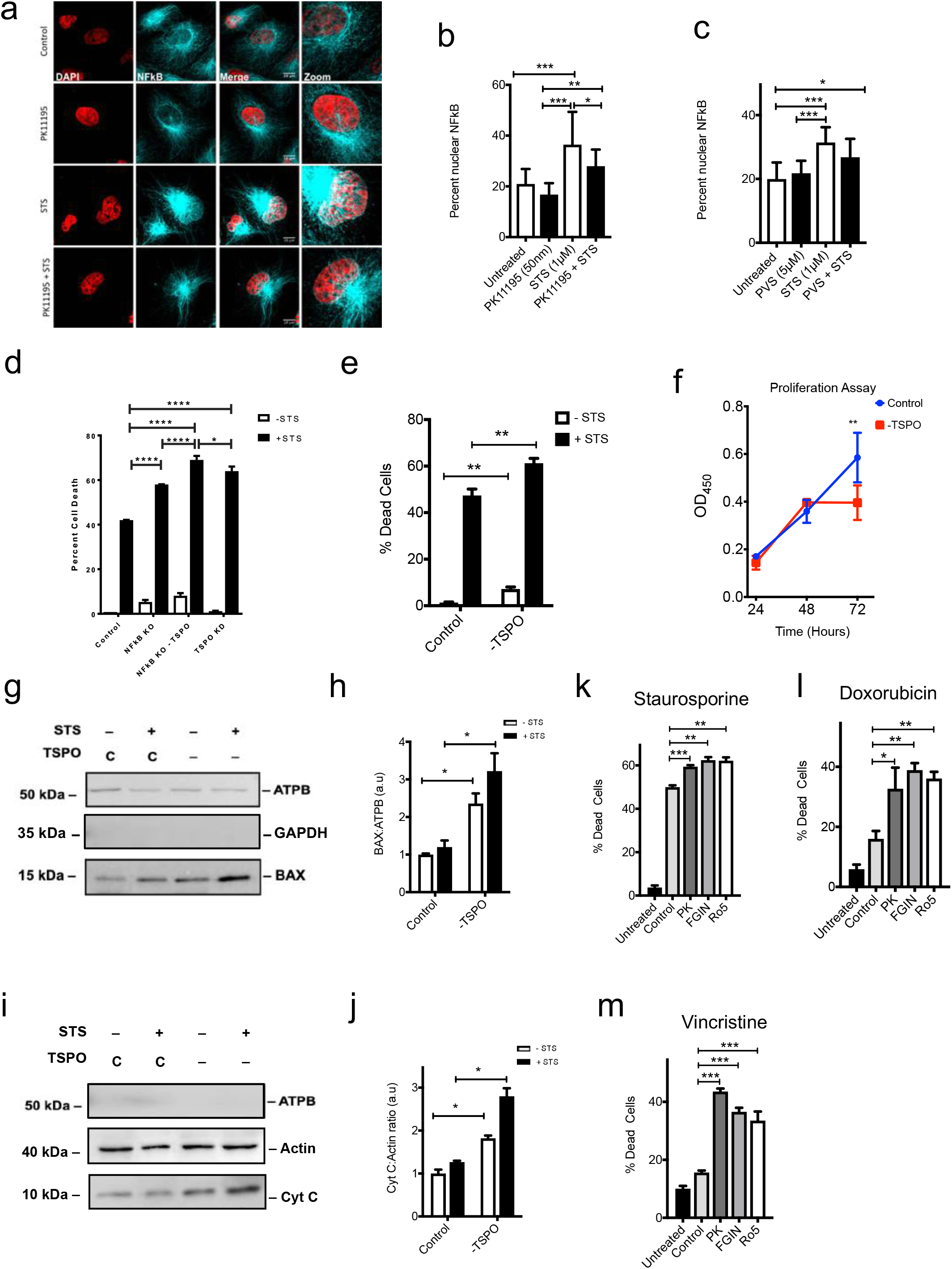
Molecular and pharmacological modulation of TSPO dictate the degree of NF-kB retro-translocation and mitochondrial commitment to apoptosis. (a) Immunocytochemistry analysis of nuclear NF-kB (Cyan) in MDA cells, untreated or exposed to STS in absence or presence of TSPO ligand PK11195. Nuclei were stained with DAPI (red). (b) Quantification of NF-κB signal in the nucleus following STS combined with PK11195. (Control 26.86±15.02; PK11195 21.26±12.24; STS 49.36±23.52; PK + STS 34.52±21.36; n=15 p<0.05, p<0.01, p<0.001) (c) Quantification of NF-κB signal in the nucleus following STS combined with Pravastatin (PVS). (Control 25.14±14.78; PVS 25.70±17.88; STS 36.20±26.64; PVS + STS 32.59±21.00; n=10 p<0.05, p<0.001) (d) Assessment of the pro-survival role of TSPO in MDA cells stably knocked-out for NF-κB, (NF-κB KO 1.3±0.69; NF-κB KO +STS 38.18±4.04; - TSPO NF-κB KO 2.88±0.07; - TSPO NF-κB KO +STS 37.1±3.45; n=3; p<0.05). (e) TUNEL assay in control and - TSPO MDA cells exposed to STS, reporting tangible differences (Control 1.37±.27; Control +STS 47.38±2.74; - TSPO 7.19 ±0.85; - TSPO + STS 61.25 ±2.06; n=3; p<0.01). (f) Proliferation assay in MDA cells highlights differences in control and - TSPO cells at 72 hours post seeding (p<0.01). (g) Western blotting analysis of Bax levels in mitochondrial fractions (marked by ATPB) of MDA cells, which reveals that the STS-induced translocation of the pro-apoptotic protein is greater in - TSPO cells. (h) Quantification of Bax band density normalised to ATPB (Control normalised 1; Control + STS 1.2 ± 0.18; - TSPO 2.35 ±.27; - TSPO + STS ±.48; n=3 p<0.05). (i) Western blotting analysis of cyt C levels in the cytosolic fractions (marked by Actin) of MDA cells, which reveals a higher degree of STS-induced cyt C translocation in - TSPO cells. (j) Quantification of cyt C band density normalised to β-actin (Control normalised 1; Control + STS 1.82±.06; - TSPO 1.27± 0.032; - TSPO + STS 2.79 ± 0.18636; n=3; p<0.05). (k) TSPO ligands PK11195, FGIN and Ro-5 exacerbate STS induced cell death in MDA-MB-231 cells (Untreated 3.71± 0.94548; Control 49.98 ±0.84; PK11195 59.43±0.68; FGIN 62.41±1.4; Ro-5 62.16±1.57, n=3, p<0.05, 0.001). (l) TSPO ligands mediate the same effect during Doxorubicin treatment (Untreated 5.9±1.52; Control 16.00±2.633; PK11195 32.7±7.03; FGIN 38.9±2.3; Ro-5 36.1±2.23; n=4; p <0.05) and (m) Vincristine induced cell death (Untreated 10.05±0.95; Control 15.61±0.67; PK11195 43.6±1.02; FGIN 36.54±1.41; Ro-5 33.53±3.13; n=3; p<0.001).

**Figure 3.**
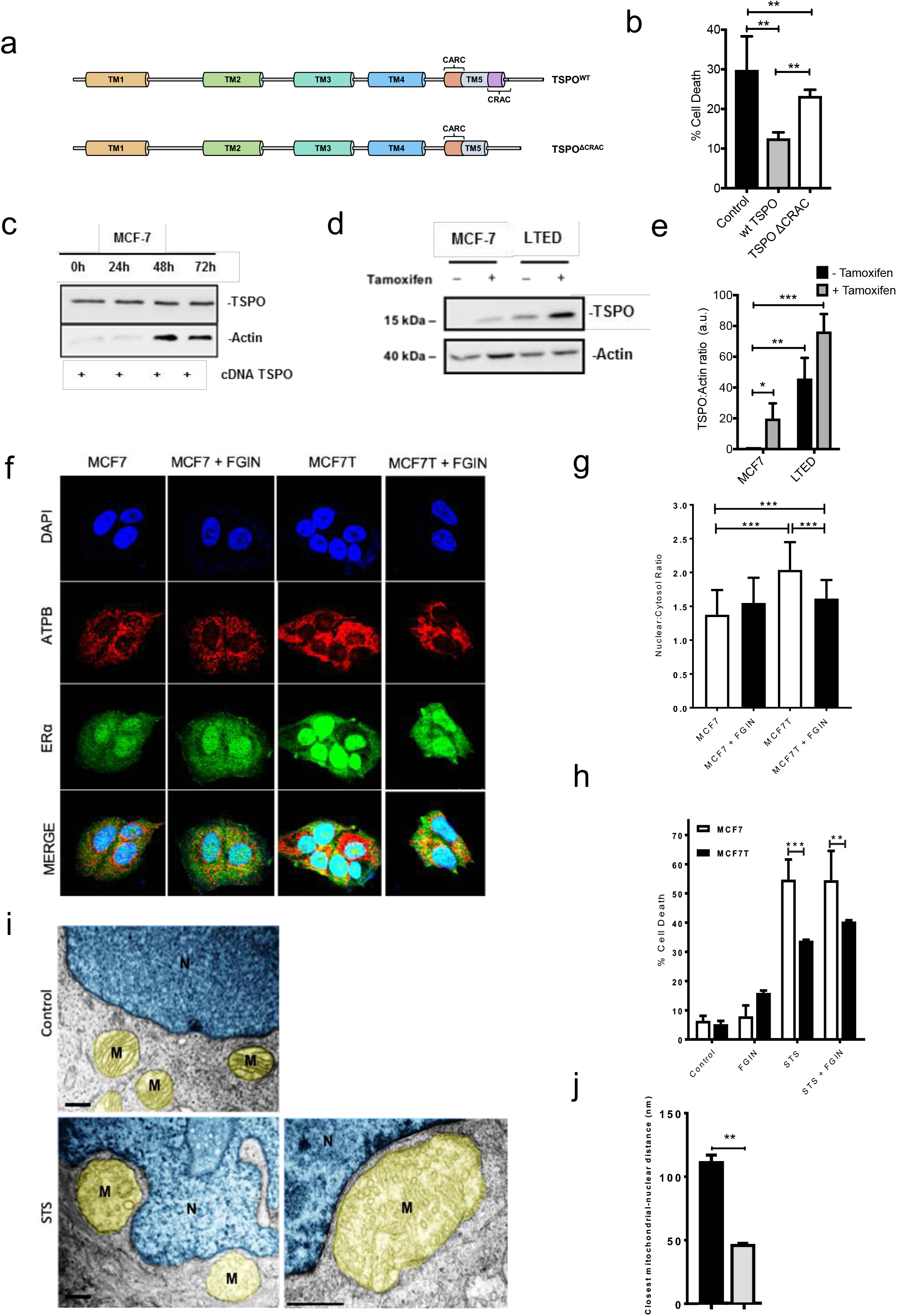
Cholesterol binding capacity is pivotal for TSPO mediated prevention of programmed cellular demise and mitochondrial associated with the nucleus. (a) Domain model of wild-type TSPO and TSPOΔCRAC construct used to knock-in the protein into MCF-7 cells. (b) Wild-type TSPO confers resistance to STS in MCF-7 cells (0.2 μM 16 h), whereas TSPOΔCRAC has a significantly lower protective effect (Control 29.86±8.48; +TSPO 15.37±4.6; +TSPOΔCRAC 2.21; n=5, p<0.005). (c) Western blot showing that expression of transgenic TSPO in MCF-7 is cells is maximal at 48 hours following transfection, and only slightly decreases after 72 hours from transfection. (d) Western blot showing that long-term oestrogen-deprived (LTED) MCF-7 cells, which are more resistant to cell death, express higher levels of TSPO when compared with parental cells, and that this expression is enhanced under tamoxifen treatment. (e) Band density analysis of control MCF-7 and LTED cells with and without Tamoxifen highlight the significant differences in TSPO expression between the two cell lines (MCF-7 Control Normalised 1; MCF-7+TAM 19.85±9.8; LTED Control 45.88±13.3; LTED + TAM 76.41±11.3; n=3; p<0.05, 0.01, 0.005) (f) Confocal imaging of MCF-7 and MCF7T cells immuno-labelled for ATPB (red) and ERα (green), with DAPI-stained nuclei. The TSPO ligand FGIN-1-27 causes cytoplasmic retention of ERα in MCF7T cells, however this effect was not observed in control MCF7 cell. (g) Quantification of nucleus-to-cytoplasm ERα ratio following treatment with the TSPO ligand FGIN-1-27. (MCF7 1.74±1.00, MCF7 + FGIN 1.92±1.18, MCF7T 2.45±1.63, MCF7T + FGIN 1.89±1.34, n=30, p<0.001). (h) MCF7T cells show resistance to STS treatment in comparison to control MCF-7. FGIN-1-27 appears to have no effect on STS-induced cell death in both cell lines (MCF7 Control 8.09±4.68, MCF7T Control 7.17±3.34, MCF7 + FGIN 11.65±4.22, MCFT + FGIN 17.33±14.58, MCF7 + STS 61.61±47.89, MCF7T + STS 34.38±33.31, MCF7 + FGIN + STS 64.56±44.49, MCF7T + FGIN + STS 41.11±39.71, n=3, p<0.001, p<0.01). (i) Representative ultrastructural images of control and STS-treated MCF-7 cells showing Remodelling of the mitochondrial network (M-yellow) and its accumulation towards the nucleus (N-blue). (j) Quantification of closest mitochondrial-nuclear distance (nm) showing reduced distance to a minimum of 50nm in STS-treated MCF-7 cells (n=5).

### Mammary cancer cells resistant to endocrine therapy upregulate TSPO whose content defines the juxtaposition of mitochondria with the nucleus

MCF-7 cells, which constitutively express lower level of TSPO, gained resistance to STS-induced apoptosis when wild-type TSPO was over-expressed. Expression of a mutant clone, devoid of the CRAC cholesterol binding domain (TSPOΔCRAC), failed instead to deliver such a protection (**Figure 3a-c**). This evidence further argued that the cholesterol-binding capacity of TSPO is crucial to drive resistance against cell death. In keeping with this, TSPO level of expression is increased in MCF-7 cells which have developed resistance to Tamoxifen (MCF7T^6^), or in which cholesterol bio-synthesis sparked following long-term oestrogen deprivation (MCF7-LTED^6,35,36^) (**Figure 3d,e**).

Treatment with the TSPO ligand FGIN-127 (FGIN), equally capable as PK11195 to impair handling of cholesterol by the protein (**Table 1**), operates as deterrent against the nuclear accumulation of the Estrogen Receptor alpha (ERα. The nuclear internalization of the latter is indeed a key event in the transition of epithelial breast tumour towards a more aggressive phenotype leading to therapeutic failure of Tamoxifen^42,43^ treatment. FGIN was beneficial in reducing ERαassociation with the nucleus (**Figure 3f,g),** as well as in increasing susceptibility to STS-induced cell death (**Figure 3h**). The co-treatment was beneficial in preventing ERαassociation with the nucleus in MCF7T than in control MCF-7 indicative of the greater availability of targets (TSPO) in this cohort of cells. In line with these observations, big-data analysis revealed that TSPO is over-expressed in cohorts of relapsed breast cancer patients, in which endocrine therapy (ET) was no longer effective and cholesterol metabolism escalated (**SFigure 3a**).

Ultrastructural imaging analysis of MCF-7 cells via transmission electron micrography (TEM) pictured the disrupted cristae and reduced mitochondrial-nuclear distance (passing from ~120nm to almost ~50nm in response to STS) (**Figure 3i,j**) which prompted us to investigate the mechanisms dictating such a relevant gain in space between mitochondria and nucleus during apoptosis.

### Mitophagy evading mitochondria, rich in TSPO, form contact sites with the nucleus in response to MRR activating conditions

Following co-immunocytochemical labelling of mitochondria with TSPO and the nuclear envelope with Lamin-B in both MCF-7 and MDA cells, we recorded that only in the latter mitochondria are constitutively present in the perinuclear region forming an association with the nucleus which is mirrored by infiltrates of TSPO-positive mitochondria (**Figure 4a,b**). This phenomenon increased in response to STS, as detailed both by immunoblotting of subcellular fractions (**Figure 4c,d**) and TEM analyses (**SFigure 2a,b**). TSPO, in response to STS, was indeed observed to be present in the portion of the cytosol occupied by mitochondria as well as in the nuclear envelope and in its core (**Figure 4c,d**). Similar dynamic redistribution of TSPO was promoted also by the mito-toxin rotenone (ROT)^44^, which is well accounted as a chemical inducer of MRR (**Figure 4c, d**).

**Figure 4.**
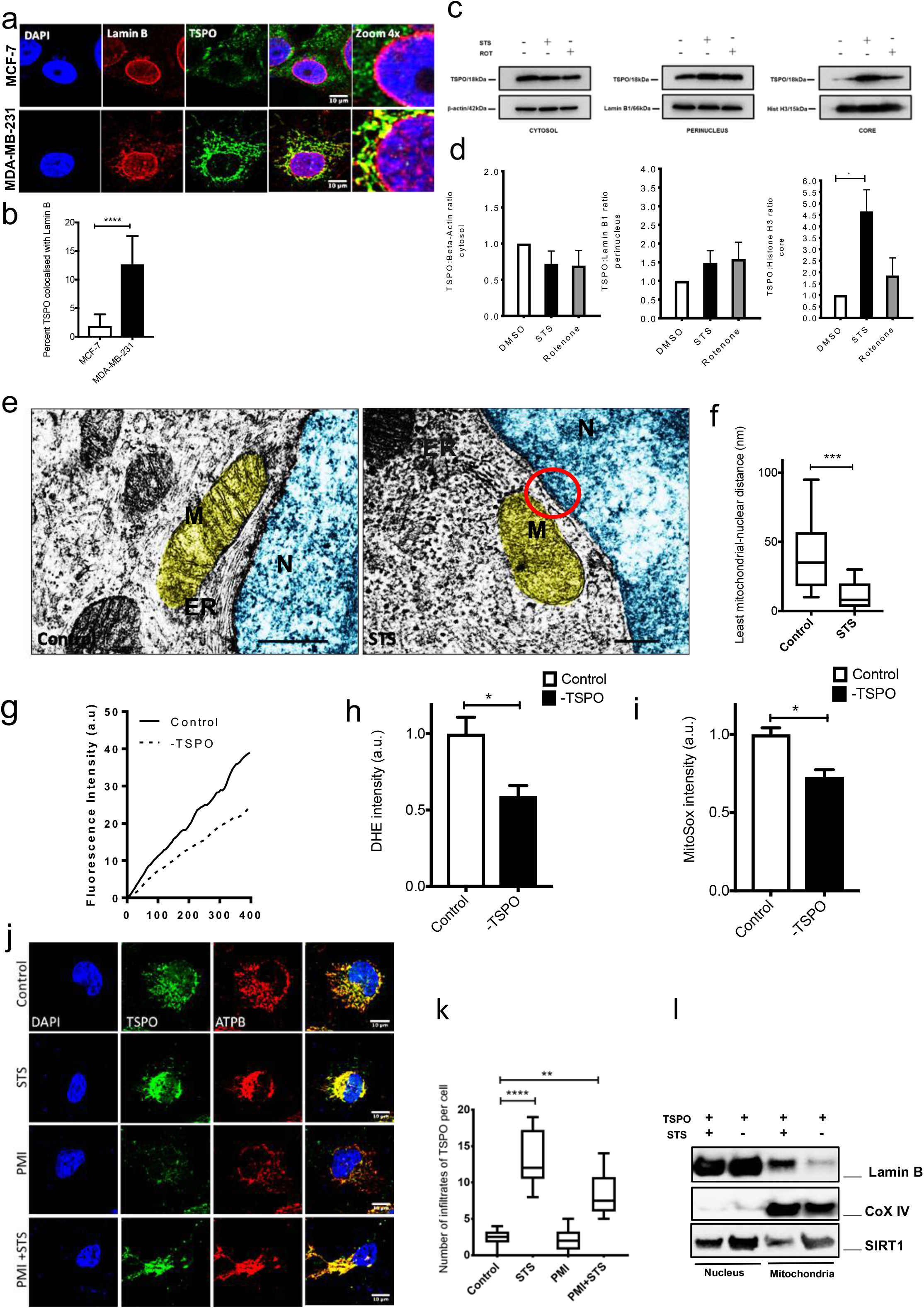
Nucleus-Associated Mitochondria (NAM) are formed during induction of mitochondrial retrograde response for which inhibition of mitophagy is required. (a) MCF-7 (top row) and MDA (bottom row) cells labelled with DAPI (blue) and immunostained TSPO (green) and lamin B1. The immunocytochemistry analysis reveals enhanced proximity of the mitochondrial network to the nucleus in MDA cells, which express greater overall amount of TSPO. (b) Co-localization of TSPO with lamin B1 in MCF-7 and MDA cells was quantified (fold-increase in MDA cells 5.40±1.5; n=3; p<0.0005) (c) Western blotting analysis of nuclear TSPO translocation following treatment with the cytotoxic compounds STS and ROT in MDA cells. From left to right, TSPO expression in the cytosol (loading control: β-actin), perinucleus (loading control: lamin B1) and nuclear core (loading control: histone H3). TSPO translocates inside the nucleus under cellular stress. (d) Densitometry analysis of TSPO levels in the different cellular fractions. Left, TSPO:β-actin ratio in the cytosolic fractions (Control DMSO normalised to 1; STS: 0.896 ± 0.543; ROT: 0.905 ± 0.487; n=3, ns). Middle, TSPO:lamin B1 ratio in the perinuclear fractions (Control DMSO normalised to 1; STS: 1.812 ± 1.162; ROT: 2.037 ± 1.137; n=3, ns). Right, TSPO:histone H3 ratio in the nuclear core fractions, showing significantly higher nuclear translocation of TSPO under stress (Control DMSO normalised to 1; STS: 5.601 ± 3.703; ROT: 2.625 ± 1.085; n=3, p<0.05). (e) TEM micrographs of control (DMSO vehicle) and STS-treated MDA cells confirming the physical interaction between mitochondria (M-yellow) and the nuclear envelope (N-blue). STS treatment causes ER displacement and the formation of contact sites between mitochondrial and nuclear membranes. (f) Quantification of minimum distance between mitochondria and the nucleus in control and STS-treated MDA cells (Control DMSO: 65.45 ± 13.89 nm; STS: 21.75 ± 1.19 nm; n=15, p<0.001). (g) Dihydroethidium (DHE)-based measurement of ROS production in control and TSPO KD MDA. DHE fluorescence intensity was plotted against time to show reduced levels of ROS production in cells devoid of TSPO (n=6). (h) Average rate of DHE fluorescence increase in control and TSPO KD MDA cells. The quantification confirms lower cytoplasmic ROS levels in - TSPO cells (Control normalised to 1;-TSPO: 0.59 ± 0.07; n=20, p<0.05). (i) Average mitochondrial accumulation of the DHE derivative MitoSox Red, reporting significantly lower mitochondrial ROS levels in - TSPO MDA cells (Control normalised to 1; - TSPO: 0.73 ± 0.04; n=20, p<0.05). (j) Confocal imaging of MDA cells exposed to STS, PMI or PMI + STS and immunolabelled for TSPO (green) and ATPB (red). DAPI was used to stain nuclei. The images reveal the collapse of the mitochondrial network, which is prevented by PMI (n=5). (k) Box plot showing the distribution of mitochondria-to-nuclear centre distances in control and STS-treated MDA cells. Under STS treatment, the distance is 4.3 μm shorter than in DMSO-vehicle control (Control DMSO: 16.17 ± 11.17 μm; STS: 11.678 ± 6.988 μm; n=9, p<0.01). (l) Western blotting analysis of the accumulation of the nuclear protein Lamin B on the mitochondria in treated (STS) and untreated MDA cells.

The ultrastructural analysis also revealed the prominent inter-organellar coupling which generated an all-encompassing structure in and around the nucleus, which we called the Nucleus-Associated Mitochondria (NAM). Means of quantification revealed the formation of actual contact sites with a statistical reduction of distance-space between mitochondria and nucleus below 30nm and observable membrane fusion (**Figure 4e,f**).

The observed mitochondrial remodelling could therefore crucially influence the signalling of diffusible molecules such as reactive oxygen species (ROS), which are acknowledged as main executors of the retrograde response^1,14^ and depending on their level of TSPO expression^16^ (**Figure 4g-i**). The NAM-induced change in space-distance would therefore favour the diffusion capacity of ROS maximizing their impact on the perinuclear and nuclear regions. Notably, the remodelling of mitochondria on the nucleus, alike the retro-translocation of NF-κB, was limited by the mitophagy-driven trimming of the network during co-treatment with PMI^31^, read by the number of TSPO infiltration in the nucleus (**Figure 4j,k**). The formation of NAM was further corroborated by assessing the level of Lamin B in purified mitochondrial fractions following STS treatment (**Figure 4l**).

### A TSPO mediated molecular complex (TSPO-ACDB3-PKA-AKAP95) tethers mitochondria to the nucleus

Formation of NAM is retained across cell types as we proved enrolling the mouse embryonic fibroblasts (MEFs), in which the expression level of TSPO defines these mito-nuclear contacts (Figure 5a-f). ‘Variance’ filter applied to confocal airy-scan images with immunofluorescence (LaminB1-ATPB) convincingly detail how the downregulation of TSPO (-TSPO) releases mitochondria from the nucleus whilst its overexpression (+TSPO) results in the coalescing of the mitochondrial network with the nuclear envelope (Figure 5a-d). These observations were corroborated at ultrastructural level via TEM analysis, which unequivocally showed the formation of contacts between mitochondria and the nucleus in the face of increased TSPO expression (Figure 5e,f). TSPO overexpressing MEFs also gained significant resistance to STS-induced cell death, which was otherwise lost when the ΔTSPO-CRAC mutant clone of the protein (unable to bind cholesterol) was expressed (Figure 5g).

**Figure 5.**
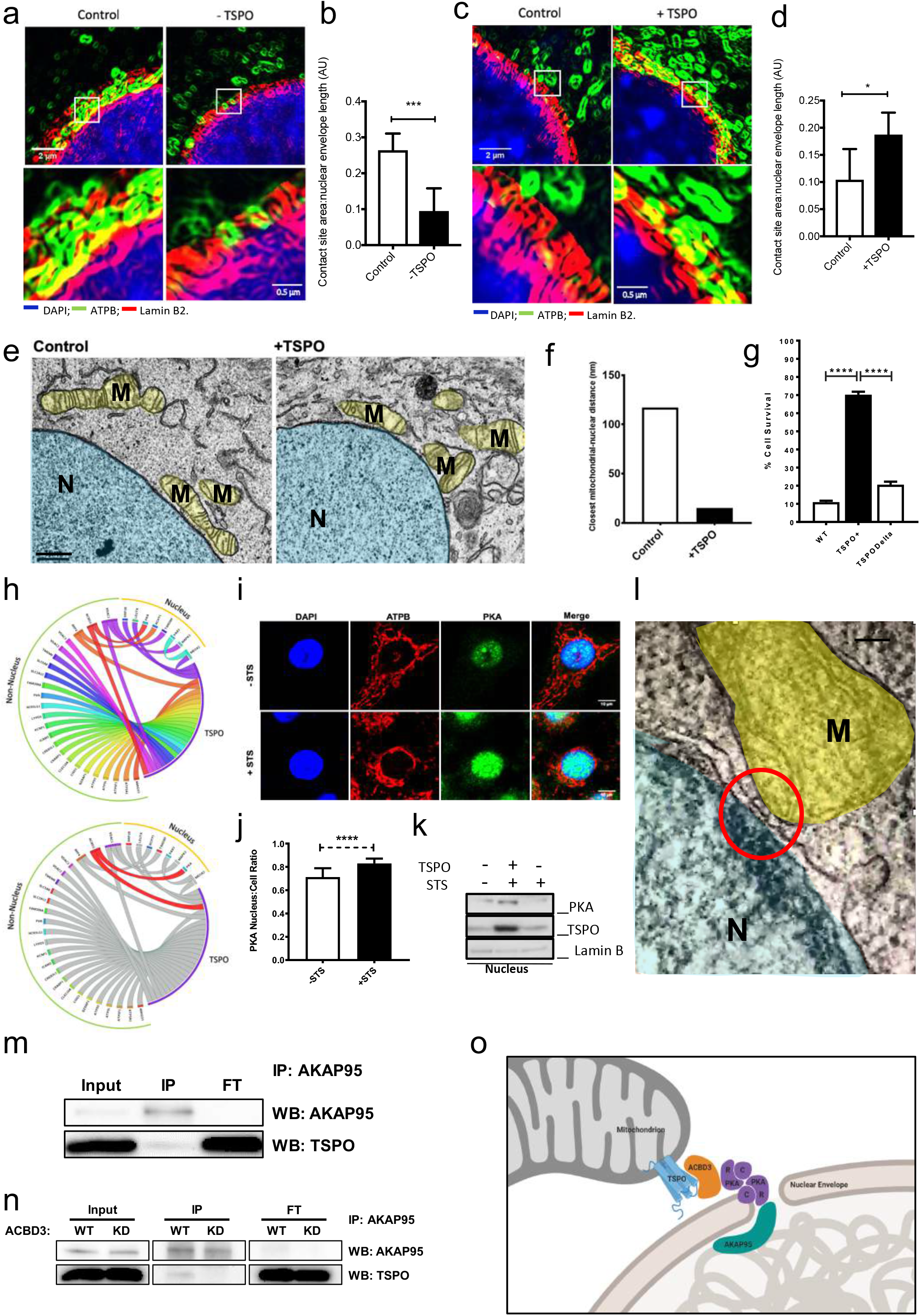
TSPO is required for the formation of mito-nuclear contact sites by forming a multimolecular tethering complex. (a) Processed Airyscan images of co-localisation between the nuclear envelope (lamin B, red) and mitochondria (ATPB, green) in scrambled siRNA control and TSPO knockdown MDA cells. (b) Quantification of contact site area in control and TSPO KD MDA cells, normalised to the nuclear envelope circumference. Knocking-down TSPO prevents mitochondria-nucleus contact site formation in MDA cells (Control: 0.3106 ± 0.213 A.U.; - TSPO: 0.15623 ± 0.03181 A.U.; n=5, p<0.01). (c) Airyscan super resolution confocal images of co-localisation between the nuclear envelope (lamin B, red) and mitochondria (ATPB, green) in control and TSPO overexpressing MEFs. (d) Quantification of contact site area normalised to the nuclear envelope circumference. Overexpressing TSPO increases mitochondrial-nuclear contact sites in comparison to control MEF cells (Control: 0.16073 ± 0.04767 A.U.; +TSPO 0.2278 ± 0.147 A.U., n=3, p<0.05). (e) Representative TEM micrograph of control and wild-type TSPO over-expressing MEFs showing mitochondrial proximity to the nucleus in the two conditions. Mitochondria are highlighted in yellow (M) and nucleus in blue (N). (f) Quantification of minimum distance between mitochondria and the nucleus in control and TSPO over-expressing MEFs (n=1). (g) STS-induced cell death assay in MEFs transfected with an empty plasmid (WT), wild-type TSPO (+TSPO) or TSPOΔCRAC (TSPOΔ). The bar chart shows that over-expressing TSPO protects from STS-mediated cell death, an effect that is lost when the CRAC domain is deleted (WT: 11.7449 ± 10.1751 %; +TSPO: 71.865 ± 68.555 %; TSPOΔCRAC: 22.243 ± 18.977 %; n=2, p<0.001). (h) Graphical representation showing nuclear and non-nuclear TSPO molecular interactors. The analysis was used to identify proteins interacting with TSPO during the formation of mitochondria-nucleus contact sites. (i) Representative TEM micrograph of a mitochondria-nucleus contact site in MEFs, suggesting the presence of a specialized tethering structure between the mitochondrial and nuclear membranes. (j) Immunofluorescence analysis of PKA (green) accumulation inside the nucleus during STS-induced MRR in MDA cells. ATPB (red) was used to stain mitochondria and DAPI (blue) to highlight the nucleus. (k) Quantification of nucleus:cytosol PKA fluorescence ratio during STS-mediated MRR. (l) Western blotting analysis of nuclear PKA and TSPO levels in MDA cells in basal conditions and after over-expression of TSPO and treatment with STS, confirming the immunocytochemistry reported in panel i. (m) Co-immunoprecipitation of AKAP95 and TSPO proteins in MDA cells. Anti-AKAP95 antibody was bound to magnetic beads and membrane were tested for AKAP95 and TSPO. 10% lysate was used as input, and 10% flow-through (FT) was added as control. (n) Co-immunoprecipitation of AKAP95 and TSPO proteins in MDA cells transfected with a scrambled siRNA (WT) or an siRNA against ACBD3 (KD). Anti-AKAP95 antibody was bound to magnetic beads and membrane were tested for both AKAP95 and TSPO. The blot shows that, when ACBD3 expression is knocked-down, the binding of TSPO to AKAP95 is prevented. (o) Diagram depicting the hypothetical anchoring platform tethering the mitochondrial membrane to the nuclear envelope. TSPO, which resides on the MOM, and nuclear matrix-localized AKAP95 can be linked together through a bridge formed by the TSPO-interactor ACBD3 and the pool of nuclear PKA.

We then attempted to dissect the mechanisms by which TSPO could mediate such a tethering role between mitochondria and nucleus hypothesising a partnering activity with other molecules. Bioinformatic analysis outlined the TSPO interactors (**Figure 5h**), and among these we teased out those with established anchoring capacity and capable of an alternative intracellular location from the mitochondria. From the analysis emerged ACBD3, which is a multi-functional interactor of TSPO and protein kinase A (PKA)^23,24^. Considering that PKA, under stress, is recruited on the nucleus and anchored via AKAP95^25,26^, we therefore hypothesized the formation of a multiprotein complex between TSPO-ACBD3-PKA-AKAP95 able to tether mitochondria with the nucleus (**Figure i**). This is evident under stress when PKA pools on the nucleus increase **(Figure 5 j-l).** To corroborate this further we run a serial of Co-IP analyses to unveil whether TSPO (the mitochondria resident end of the complex) and AKAP-95 (the nucleus resident end of the complex) interact. In MDA cells in which the expression of TSPO is endogenously high the interaction between TSPO and AKAP95 was proved as well as its regulation by ACBD3 **(Figure 5 m,n).** Thus, knocking down ACBD3 (-ACBD3) does succeed in reducing the association between mitochondria and nucleus (**SFigure 4 a,b)**. Based on this a topology of the tethering complex is therefore proposed in **Figure 5 o.**

The distinctive molecular nature of NAM from Mitochondria-associated ER membranes (MAM), archetypes of the mitochondrial interaction with other organelles, was subsequently tested by assessing their degree following modulation of the integral membrane protein VAPB5 and Mitofusin 2^45^. Removal of either of them did not affect the connection between mitochondria and the nucleus (**SFigure 3j,m and SFigure 4c,d**).

### NAM exploit TSPO to redistribute cholesterol into the nucleus and promote NF-κB activity

The formation of NAM via the upregulation of TSPO is promoted by the oxidized forms of cholesterol, as demonstrated by the exposure of MDA cells to 7-ketocholesterol (7-KC) (**Figure 6a-e**), which contributes to the etiopathology of several conditions including those characterized by malignant proliferation^46,47^.

**Figure 6:**
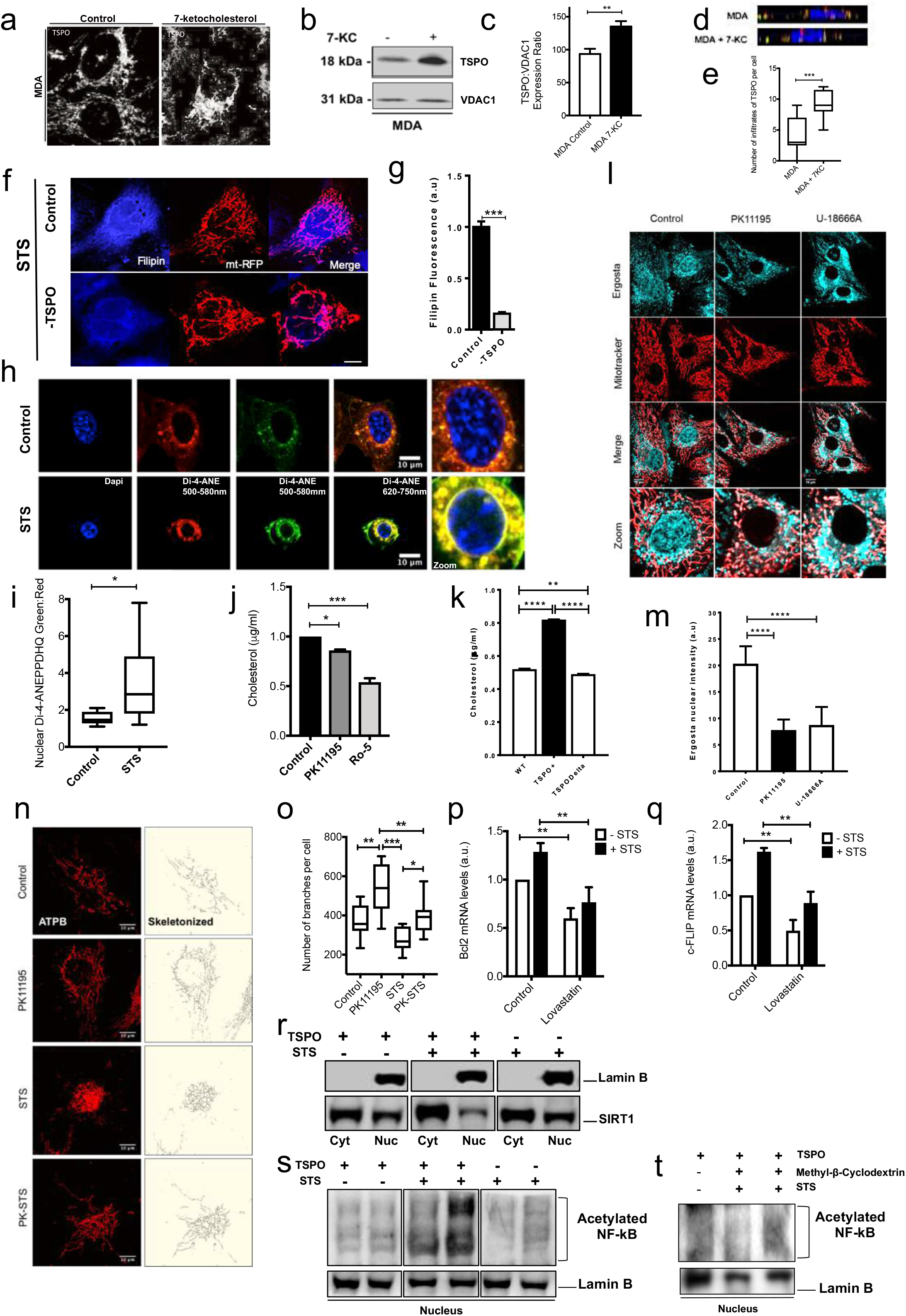
Redistribution of cholesterol in the nucleus aids NF-κB activity, is secondary to NAM formation and requires functional TSPO. (a) Representative confocal images of MDA cells, control and incubated with the toxic oxysterol 7-ketocholesterol (7-KC), immunostained for TSPO, showing upregulation of the protein after treatment. (b) Western blot confirming TSPO upregulation in MDA cells following treatment with 7-KC. (c) Densitometry analysis of TSPO protein levels (normalized to VDAC) in MDA cells untreated or incubated with 7-KC, showing statistically significant upregulation of TSPO induced by 7-KC (Control vehicle: 94.861 ± 6.5, 7-KC: 136.59 ± 6.76; n=3; p<0.01). (d) Prototypical orthogonal projection of z-stack confocal images of MDA cells, untreated or incubated with 7-KC, immunostained for TSPO (red) and lamin B (green). The number of TSPO rich mitochondria infiltrating the nucleus (stained in blue with DAPI) in 7-KC treated MDA-MB-231 cells compared to untreated ones. (e) The total number of TSPO-positive infiltrates per cell was quantified in both control and 7-KC-treated MDA cells (Control vehicle: 4.22 ± 2.9; 7-KC: 12.34±2.3; n=9, p<0.005). (f) Confocal images of MDA-MB-231 cells modulated for TSPO expression, transfected with mitochondria-targeted RFP (mt-RFP, red) and stained with the cholesterol marker filipin (cyan). The images show reduced cholesterol in the perinuclear region in - TSPO cells (bottom row). (g) Quantification of nuclear filipin staining in MDA cells, highlighting that - TSPO cells have reduced accumulation of the dye when compared with control. (Control: 1.04 ± 0.009; - TSPO: 0.2 ± 0.001; n=8, p<0.001) (h) Confocal images of MDA cells following STS treatment stained with the fluorescent probe Di-4-ANEPPHQ (ANE), which senses lipid rigidity and was used as readout of nuclear membrane stiffness and cholesterol content. Under STS treatment, the nuclear envelope becomes more ordered, as indicated by the increase in ANE green fluorescence, confirming an increase in cholesterol-enriched lipid domains. (i) Box plot of ANE green:red fluorescence ratio in the nucleus of MDA cells, showing a statistically significant blue-shift of the dye following STS treatment. (j) Bar chart reporting the nuclear cholesterol content in control MDA cells and after incubation with the two TSPO ligands PK11195 and Ro-5 (Control normalised: 1; PK11195: 0.86 ± 0.008 μg/ml; Ro-5: 0.54 ± 0.039 μg/ml; n=3, p<0.001). (k) Evaluation of nuclear cholesterol levels in MEFs expressing either wild-type TSPO (TSPO+) or TSPOΔCRAC (TSPODelta). Data show that the cholesterol-binding capacity of TSPO is required for the nuclear accumulation of cholesterol. (l) Confocal live-cell imaging analysis of the intracellular trafficking of the cholesterol analogue *ergosta*-5,7,9(11),22-tetraen-3β-ol (ergosta, cyan). CF35 canine mammary cells were co-stained with Mitotracker Red CMXRos (red) to visualize the mitochondrial network. The TSPO ligand PK11195 prevents the uptake of ergosta into the nucleus, similarly to the cholesterol trafficking inhibitor U-18666A. (m) Quantification of ergosta nuclear intensity in CF35 cells following treatment with either PK11195 or U-18666A (Control vehicle: 23.629 ± 17.031; PK11195 9.784 ± 5.814; U-18666A 12.164 ± 5.362; n=25, p<0.001). (n) Representative confocal images of MDA cells immuno-labelled for ATPB (red, left), with corresponding skeletonized representation (right) of mitochondrial network in cells treated with PK11195, STS or a combination of the PK11195 and STS. (o) Box plot of mitochondrial branches per cell reveals that PK11195 treated cells have far more arborizations, which also are retained during STS treatment (Control vehicle: 371.6 ± 82.3 branches/cell; PK11195 532 ± 123 branches cell; STS: 275 ± 62.9 branches/cell; PK11195+STS: 390 ± 86.5 branches/cell; n=10, p<0.01, 0.005). (p) qPCR analysis of the ability of the cholesterol synthesis inhibitor Lovastatin to reduce Bcl2 expression after treatment with STS in MDA cells; data show a statistically significant inhibition of Bcl2 transcription under Lovastatin treatment (Control vehicle normalised to 1; control + STS: 1.08 ± 0.04; Lovastatin vehicle: 0.6 ± 0.18; Lovastatin + STS 0.77 ± 0.2; n=3, p<0.01). (q) Bar chart reporting qPCR analysis of c-FLIP mRNA levels in the presence of Lovastatin in both control conditions and STS treated MDA cells (Control vehicle normalised to 1; control + STS 1.49±0.32; lovastatin.052±0.1; lovastatin +STS.85±0.10; n=3; p<0.01). (r) TSPO favours repression of Sirtuin1 accumulation in the nucleus during MRR. (s) NF-kB de-acetylation raises during MRR and is prevented by TSPO repression. (t) Methyl-β-cyclodextrin by depleting cholesterol blocks NF-kB de-acetylation during STS.

In response to 7-KC, the identical pattern of TSPO expression is mirrored by other components of the steroidogenic protein complex-transduceome^19^, ATAD3 and StAR, and the oxysterol receptor LXR-β^48^ (**SFigure 2f-j**). This outlines an interplay between mitochondrial handling of cholesterol, its metabolism and the tonicity of the MRR, which we investigated by monitoring the redistribution of cholesterol after remodelling of the mitochondrial network in response to STS treatment. In MDA cells expressing the red variant of the green fluorescent protein (mt-RFP), immuno-labelled with Lamin-B1 to stain the nuclear envelope, we imaged the cholesterol pools via the fluorescent compound filipin, which depicted the localization of the lipid into foci close to and around the nucleus following STS treatment. These foci disappeared in -TSPO MDA cells (**Figure 6f,g**).

The stress induced retro-trafficking of cholesterol was confirmed by imaging the lipid enrichment of the nuclear membranes in MDA cells with the fluorescent probe Di-4-ANEPPDHQ (**Figure 6h, i**). In the same conditions, the efflux of cholesterol from the nucleus could be observed during cell treatment with TSPO ligands (**Figure 6j**), as well its marked influx when the protein is overexpressed (**Figure 6k**). The pattern of cholesterol was also assessed by live imaging using dehydroergosterol [ergosta-5,7,9(11),22-tetraen-3β-ol, DHE]^49^, via which we observed the inhibition of nuclear redistribution of cholesterol by STS when cells are co-treated with the TSPO ligand PK11195^50^ or the intracellular cholesterol transport inhibitor U1866A as positive control (**Figure 6l,m)**. Notably, the pharmacological targeting of TSPO did also prevent remodelling of mitochondria (**Figure 6n,o**).

The inter-dependence between cholesterol accumulation in the nucleus and transcriptional activity of NF-κB was finally confirmed by monitoring the expression levels of the pro-survival genes (c-FLIP, Bcl-2), both of which were reduced by the cholesterol synthesis inhibitor Lovastatin (**Figure 6p,q**), likewise it is in -TSPO cells (**Figure 1n,o**). Furthermore we gained evidences that the increased level of NF-κB de-acetylation hence activation during STS treatment associates with the reduced nuclear accumulation of the nutrient-sensing deacetylase Sirtuin-1 (SIRT-1)^51,52^ dependent which is otherwise favoured by the repression of TSPO level (**Figure 6r,s**). Depletion of cellular cholesterol via Methyl-β-cyclodextrin mimics similar protective effect on de-acetylation of the NF-κB (**Figure 6 t**).

The final graphical model in **Figure 7** summarises the key steps of this pathway in which association of mitochondria with the nucleus and redistribution of cholesterol regulate MRR activation and execution.

**Figure 7.**
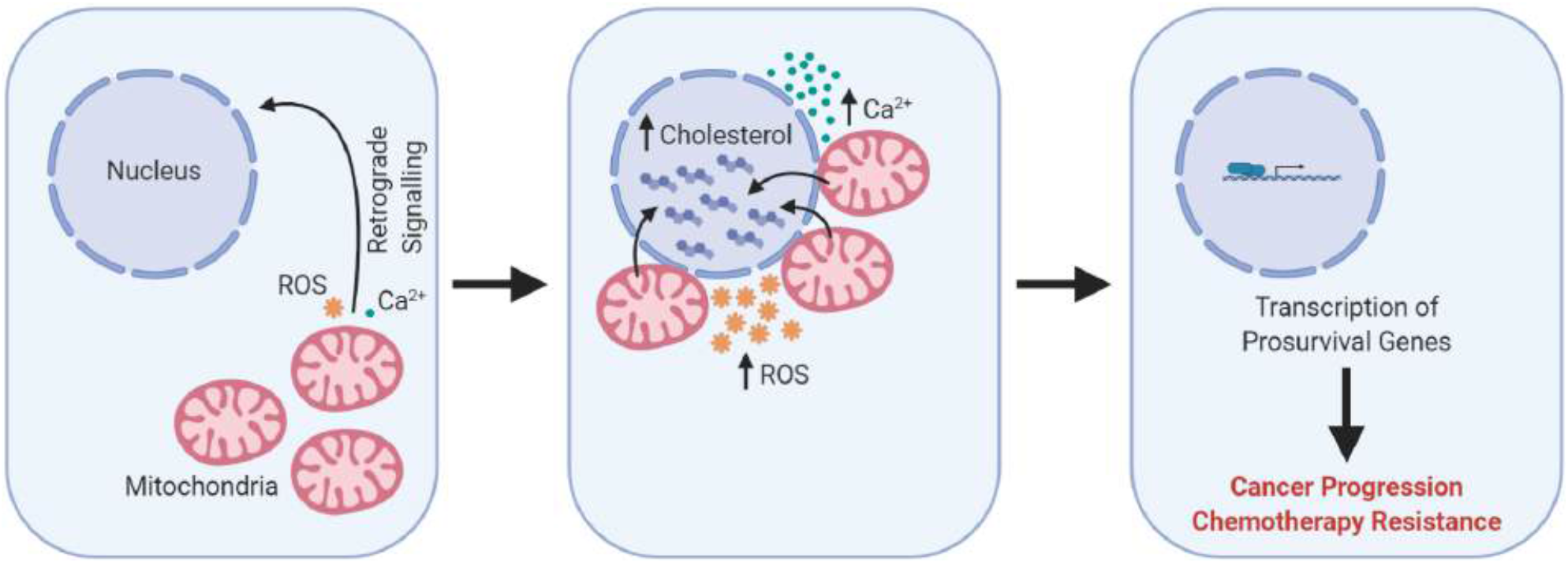
Graphical representation of the working model on the formation of contact sites between mitochondria and nucleus (called NAM) during the execution of the mitochondrial retrograde response.

## Discussion

In this work, we demonstrate that mitochondria can establish points of contact with the nucleus to favour spatially confined biochemical events that promote nuclear stabilization of pro-survival transcriptional factors such as NF-κB. We called these newly identified contact sites Nucleus-Associated Mitochondria (NAM). We also report that cholesterol works as an intermediate or a facilitator of the mitochondria-to-nucleus retro-communication together with ROS and Ca^2+^, which have been acknowledged as the prevalent dynamic signalling molecules between the two organelles^1,2^. The accumulation of cholesterol in the nuclear envelope emerges as regulator of cellular reprogramming towards stress resistance and survival (**Figure 6f-m**). Even though cholesterol is known to affect transcription of genes^48^, it is here for the first time reported to be trafficked into hot-spots at the interface between mitochondria and the nucleus to promote their communication, thus resembling other platforms for the exchange of information such as those between the ER and mitochondria (ER-MAMs)^53^ which we prove to be molecularly distinct from the NAM (**SFigure 4**).

Following mitochondrial re-organization, TSPO may therefore deliver its lipid cargo, cholesterol, into the nuclear core of the cell where the concurrence of ROS may lead to auto-oxidative products of cholesterols (oxysterols^47^). The pattern of elements of the transduceome-complex^31^, which all converge into the nucleus along with TSPO strongly argues for this (**SFigure 2e-j**). We speculate that the enrichment of the nuclear envelope with lipids such as cholesterol, inducing a change in the physical properties **(Figure 6h)**, represents a crucial step in the altered transcriptional profile which enables aggressive cancer cells to differentiate from their less persistent counterparts. Here we unveiled that cholesterol is pivotal for the efficiency of the retrograde response as the degree of NF-κB acetylation is affected by its redistribution in the nucleus **(Figure 6r-t)**, which represses the NAD-dependent deacetylase sirtuin-1 as previously proposed^51,52^.

While the observation that the mitochondrial interactome is crucial to cellular health is not new^54^, the fine regulation of mitochondrial juxtaposition on the nucleus is. We have here detailed that TSPO is required for this inter-organellar interaction, and in turn, its ligands work as efficient pharmacological modulators of the coupling (**Figure 6l,m**). TSPO is therefore required for the physical interaction between the two membranes, and we propose this to be mediated by a complex of proteins formed by the scaffolding protein ACBD3 which bridges the PKA complexed with AKAP-95 on the mitochondria. This complex is pictured via Tomograph analysis (**Figure 5 I and SFigure 4 e**) and graphically outlined in **Figure 5o**. Even though we cannot rule out that VDAC is required to ensure the right positioning of the OMM to tether with the nucleus we envisage a relay of proteins which interplay during stress such as TSPO, ACBD3 and PKA^23,24^.

TSPO is an ancient protein^55^ and its key part in the physical fine-tuning between two genomes may be an evolutional adaptation taking place via the exploitation of a lipid binding protein to fulfil both a physical and functional interaction.

In cellular models of malignant lesions of the breast, such as MDA cells, NAM are indeed prominent, far more than in the less aggressive counterpart MCF-7 cells. In these, STS treatment is nonetheless able to promote a statistically significant reduction in the distance between mitochondria and the nucleus (**Figure 3i,j**), even though still not within the range of contacts as it does in MDA cells, in which TSPO level is very high (**Figure 4a,b**). This steady-state formation of NAM may be an ‘already established’ communication platform in these cells which is exacerbated by cholesterol dysregulation and therapy treatment.

NAM will in turn increase the exposure of the nucleus to hydrogen peroxide (H_2_O_2_), superoxide (O_2_-) as well as hydroxyl radical (OH^−^) generated in excess by defective mitochondria overexpressing TSPO. The nucleus is usually out of reach of mitochondrial O_2_- and OH^−^, which have very low estimated diffusion co-efficients^56^. The formation of NAM would bring mitochondria in a closer range to the nuclear DNA facilitating the exposure of reactive species of the oxygen to the nuclear DNA^57^ which maximize NF-κB activation.

NAM may therefore represent both a defensive mechanism in response to apoptotic cues in the short term, as well as a detrimental coupling that compromises DNA integrity in the long one, leading to genomic alterations. This ties well into the observations that cytoskeletal reorganization (which we have not here investigated) and cellular morphology are key factors in cellular resistance to apoptosis as well as in regulating the NF-κB signalling^58,59^. Equally, nuclear pleomorphism of tumours, proposed as a biomarker of malignancy for breast cancer^60^, and the associated somatic mutations could be the consequence of the continuous exploitation of this localised, but nuclear-focused, stress signalling.

Mitochondrial repositioning is nonetheless secondary to inefficient mitochondrial quality control for impaired mitophagy^61^, which causes the accumulation of inefficient organelles and their bypassed respiratory products. TSPO suppresses PINK1-PARKIN-driven mitophagy by preventing the ubiquitination of mitochondrial proteins^16^. TSPO rich mitochondria are therefore prone to repositioning on the nucleus, as they can systematically evade control by autophagy. The previously described de-ubiquitinating properties of TSPO^16^, lead us to speculate that trafficking of TSPO onto the nuclear envelope could replicate a similar outcome in the nucleus by preventing nucleophagy^62^ of which the undulated nuclear envelope observed ultra-structurally could be the consequence (**SFigure 2a**). The alteration in lipid composition of the membranes to which TSPO localises (mitochondria) or re-localises (nucleus) could also be instrumental for this as well as to the altered organellar life cycle and protein interactome (**Figure 5h**). Mitochondrial network remodelling, cholesterol sequestration on the perinuclear region and cell death evasion are indeed avoidable by the downregulation of TSPO, its pharmacological inhibition as well as the pharmacological activation of mitophagy. Furthermore, TSPO is overexpressed in mammary gland cell lines that have developed resistance to ET thus conferring resistance to those who might instead be susceptible to the treatment (**Figure 4 d-g**). Overexpression of the ΔTSPO-CRAC mutant, which fails binding of cholesterol by TSPO, is unable to replicate the same effect (**Figure 5g**). These experiments propose therefore a critical role for cholesterol in the resistance behaviour primed by the MRR, which is executed by physical re-organization of the mitochondrial network on the nucleus to via specific contacts, which have been here reported for the first time.

## Author Contribution

M.C. conceived, designed and coordinated the project together with R.D, and D.A.E. M.C. R.D., L.H., D.A.E, J.C., D.F., M. S. A. performed the experiments and ran the analysis. R.F., G.V and L.A performed the EM analysis whilst G. T., M.M. the immunohistochemistry analysis on which guidance was obtained by K.S and V.Z., R.B. and G.S. worked out the bioinformatics data. R.A. run and analysed the biochemical experiments on Sirtuin-1. All authors have critically reviewed the manuscript and advised accordingly.

## Acknowledgments

The research activities lead by M.C. are supported by the following funders, who are gratefully acknowledged: Biotechnology and Biological Sciences Research Council [grant numbers BB/M010384/1 and BB/N007042/1]; The European Research Council COG 2018 - 819600_FIRM; Bloomsbury Colleges Consortium PhD Studentship Scheme; The Petplan Charitable Trust; Umberto Veronesi Foundation; Marie Curie Actions and LAM-Bighi Grant Initiative. FIRB-Research Grant Consolidator Grant 2 [grant number: RBFR13P392], Italian Ministry of Health [IFO14/01/R/52].

## Methods

### Immunohistochemistry/Immunocytochemistry

Immunofluorescence: Cells were fixed in 4% PFA (10 mins, RT) followed by 3 five-minute washes in PBS. Permeabilization was performed with 0.5% Triton-X in PBS (10 mins, RT) followed by washing. Blocking was carried out for 1h at RT in 10% Goat Serum and 3% BSA in PBS. Primary antibody incubations were conducted overnight for 16 h at 4°C in blocking solution as described. After a further wash step, secondary antibodies were incubated for 1 h in blocking solution, before a final wash step. Cells were then mounted on slides with DAPI mounting medium (Abcam, ab104139). Cells were stained with the following primary antibodies: ATPB (Abcam ab14730) 1:400, Lamin B2 (Abcam ab8983) 1:500; TSPO (Abcam, ab109497) 1:200, NF-kB (AbCam, ab16502) 1:500, 1:1000 ATAD3 (Gift from Dr Ian Holt, Biodonista, 1:400 LXRβ (Abcam ab28479), 1:500 STAR (AbCam ab58013); and the following secondary antibodies: α-mouse Alexa 555 (Life Technologies, A21424) 1:1000; α-rabbit Alexa 488 (Life Technologies, A11008) 1:1000.

Single dye immunofluorescence was used to stain paraffin sections of human mammary tissues with different TSPO antibody (ab109497) / NF-kB (ab32360). The binding of mouse primary antibodies was detected using Alexa Fluor 488 (or 594) fluorochrome-conjugated goat anti-mouse IgG. Tissue sections were mounted with Floroshield Mounting Medium with DAPI to show

### Cell Culture

MDA-MB-231 and MCF7 human breast cancer cell lines, CF41 canine mammary line, K248P feline mammary line, WT and Mitofusin 2^−/−^ cell lines were all maintained at 37°C under humidified conditions and 5% CO_2_ and grown in Dulbecco’s modified Eagle medium (Life Technologies, 41966-052) supplemented with 10% foetal bovine serum (Life Technologies, 10082-147), 100 U/mL penicillin, and 100 mg/mL streptomycin (Life Technologies, 15140-122). K248P cells were additionally supplemented with 10 μg/ml insulin (sigma-aldrich). Cells were transiently transfected with genes of interest or siRNA using a standard Ca^2+^ phosphate method as described previously^63^ or using manufacturers’ instructions for Lipofectamine 3000 (Thermofisher 18324010). Mnf2KO MEFs were obtained from the Laboratory of Prof. Luca Scorrano (University of Padua, Italy).

### Cell Fractionation

Cells were lysed in cold isotonic buffer (250 mM Sucrose, 10 mM KCl, 1.5 mM MgCl_2_, 1 mM EDTA, 1 mM EGTA, 20 mM HEPES, pH 7.4) containing protease inhibitor cocktail (Roche, 05892791001) by passing through a 26-gauge needle 10 times using a 1 ml syringe, followed by 20 minutes incubation on ice. Nuclei were separated by centrifugation at 800g for 5 minutes at 4°C. For nuclear fractions, pellets were washed once in isotonic buffer followed by centrifugation 800g for 10 minutes at 4°C. Pellets were suspended in standard lysis buffer (150 mM NaCl, 1% v/v Triton X-100, 20 mM Tris pH 7.4) plus 10% glycerol and 0.1% SDS and sonicated for 5 seconds to dissolve pellet. For mitochondrial fractions, supernatants were transferred to fresh tubes and centrifuged at 10000g for 10 minutes at 4°C, Subsequent supernatants were collected as the cytosolic fractions while mitochondrial pellets were washed once in cold isotonic buffer then centrifuged at 10000g for 10 minutes at 4°C. Finally, mitochondrial pellets were lysed in lysis for 30 minutes on ice.

### Western Blot

Sample proteins were quantified using a BCA protein assay kit (Fisher Scientific, 13276818). Equal amounts of protein (10-30 μg for whole cell lysates/cytosolic fractions; 10 μg for mitochondrial. Nuclear fractions) were resolved on 12% SDS-PAGE gels and transferred to nitrocellulose membranes (Fisher Scientific, 10339574). The membranes were blocked in 3% non-fat dry milk in TBST (50 mM Tris, 150 mM NaCl, 0.05% Tween 20, pH 7.5) for 1 h then incubated with the appropriate diluted primary antibody at 4°C overnight: LC3 (Abcam, ab48394) 1:1000; TSPO (Abcam, ab109497) 1:5000; Actin (ab8266). 1:2000; ATPB (Abcam, ab14730) 1:5000, BAX (Abcam, ab32503) 1:1000; CytC (ab13575) 1:1000; Histone H3 (ab8580) 1:1000 NF-kB (ab32360 1:1000; Lamin B1 (ab16048) 1:1000; MTCO1 (ab14705) 1:1000. Membranes were washed in TBST (3 × 15 mins at RT) and then incubated with corresponding peroxidase-conjugated secondary antibodies (Dako, P0447, P0448) for 1h at RT. After further washing in TBST, blots were developed using an ECL Plus on western blotting detection kit (Fisher Scientific, 12316992). Immunoreactive bands were analyzed by performing densitometry with ImageJ software.

### Confocal Imaging/ImageJ

Fluorescent labeling was observed through a LSM 5 Pascal confocal microscope (Zeiss, Oberkochen, Germany) and images were recorded with the Pascal software (Zeiss). All image analysis was done using Fiji (ImageJ; NCBI, USA) and corresponding plug-ins. All staining was checked for non-specific antibody labeling using control samples without primary antibody. None of the controls showed any signs of nonspecific fluorescence. For ICC and IHC imaging, single optical sections were analyzed. The infiltrate analysis was done using orthogonal rendering 3D optical stacks and counting the number of infiltrates per cell nucleus. The mitochondrial network analysis was performed using the ‘skeletonize’ plugin. Higher resolution airyscan processed images were acquired using an LSM 880 Confocal with Airyscan (Carl Zeiss, Jena, Germany) system with GaAsP detectors and a module for airyscan imaging. In Airyscan modes, a 63x Plan Apochromat (1.4 NA) oil objective was used. Confocal imaging was sequential for different fluorophore channels to obtain a series of axial images. Images of cells were optimized for visualization by manually adjusting the minimum and maximum intensity values of the histogram in the Zen software (Carl Zeiss). Images were analyzed using the variance filter and quantified by co-localization of fluorophores.

### Cell Proliferation assays

Cell proliferation assay were performed using the WST-1 Cell Proliferation Assay Kit (ab65473) and assessed on a standard plate reader (Tecan SUNRISE)

### Cell Death Assays

Cells were fixed in 4% PFA for 10 mins at, RT followed by 3 five-minute washes in PBS. Permeabilization was performed with 0.2% Triton-X100 in PBS for 10 mins at RT followed by washing. Death assays TUNEL Assay Kit - *in situ* Direct DNA Fragmentation (ab66108) as per the manufacturer’s instructions. Alternatively, cells were live stained with Hoechst 33342 (Sigma H6024) and Propidium iodide (Sigma 25535) to mark apoptosing cells and imaged on an inverted fluorescence microscope.

### Fluorescence imaging

Cells were incubated with 5 μM dihydroethidium (DHE, Life Technologies, D-1168) or 5 μM MitoSOX (Life Technologies, M-36008) in recording media (125 mM NaCl, 5 mM KCl, 1 mM NaH_2_PO_4_, 20 mM HEPES, 5.5 mM glucose, 5 mM NaHCO_3_, and 1 mM CaCl_2_, pH 7.4) for 30 minutes at 37 ⁰C. Cells were washed once in recording medium then transferred to a Leica SP-5 confocal microscope (63X oil objective lens) for imaging and fluorescence intensity was measured through continuous recording for at least 10 mins. Settings were kept constant between experiments. Mitochondrial ROIs were selected, and the corresponding fluorescence intensities calculated. For the transport of cholesterol assay, Ergosta (dehydroergosterol-ergosta-5,7,9(11),22-tetraen-3β-ol) was prepared and loaded as previously described^64^. Briefly, Ergosta was added to an aqueous solution of MβCD (3mM Ergosta and 30mM MβCD). This mixture was overlaid with nitrogen, continuously vortexed under light protection for 24 h at room temperature and filtered through a 0.2 μm filter to remove insoluble material and Ergosta crystals. Then, 20 μg of DHE was added to the cells in the form of DHE-MβCD complexes and allowed to incubate for 90 minutes at room temperature in PBS. Prior to imaging, cells were washed three times with culture media before being incubated with 20 nM MitoTracker™ Red CMXRos. Cell were washed and then imaged.

### Statistical analysis

Data are presented as mean ± standard deviation of the mean. One-way analysis of variance (ANOVA) was used in multiple group comparisons with Bonferroni’s *post hoc* test to compare two data sets within the group and a *p* value less than 0.05 was considered significant. All analyses were performed in Microsoft Office Excel 2010 and GraphPad Prism 7. *p = <0.05, **p = <0.01, ***p = <0.001

### Preparation of cells for Electron Microscopy (TEM and Immunogold TEM)

For TEM analysis, MDA-MB-231 cells (control and treated with 0.5 μm for 16 h) were fixed at room temperature with 2.5% glutaraldehyde (v/v) in 0.1 M sodium cacodylate buffer (pH 7.4) for 3 h. The cells were pelleted by centrifugation, washed in buffer and post-fixed for 1 h with 1% osmium tetroxide in 0.1 M sodium cacodylate buffer. Samples were thoroughly rinsed, dehydrated in a graded series of ethanols and embedded in epoxy resin (TAAB 812). Ultrathin sections (70-90 nm) were cut using a Leica UC7 ultramicrotome mounted on 150 mesh copper grids and contrasted using Uranyless (TAAB) and 3% Reynolds Lead citrate (TAAB). Sections were examined at 120kV on a JEOL JEM-1400Plus TEM fitted with a Ruby digital camera (2kx2k). For TEM immunogold labelling, cell samples were fixed with 4% (w/v) paraformaldehyde, 0.1% (v/v) glutaraldehyde in 0.1M phosphate buffer (pH 7.4) for 4 h at room temperature and spun down on 20% gelatine. Gelatine cell pellets were cryoprotected by incubating in 2.3M sucrose overnight at 4°C. Gelatine blocks containing cells were cut further into 1-2mm cubes, mounted on aluminium pins and cryofixed by plunging into liquid nitrogen. Samples were stored in liquid nitrogen prior to cryosectioning. Ultrathin sections (70-90 nm thick) were cut using a Leica EM FC6 cryo-ultramicrotome and mounted on pioloform film-supported nickel grids according to the Tokuyasu method (1). Sections were immunolabeled using anti-TSPO (Abcam, ab109497) (1:200) followed by a 12nm-colloidal gold anti-rabbit secondary antibody (Jackson ImmunoResearch) (1:40). Grids were examined at 120 kV on a JEOL JEM-1400Plus TEM fitted with a Ruby digital camera (2k×2k).

