## Supplementary Figures for "Mitochondria form contact sites with the nucleus to couple pro-survival retrograde response"

### Supplementary Figure 1

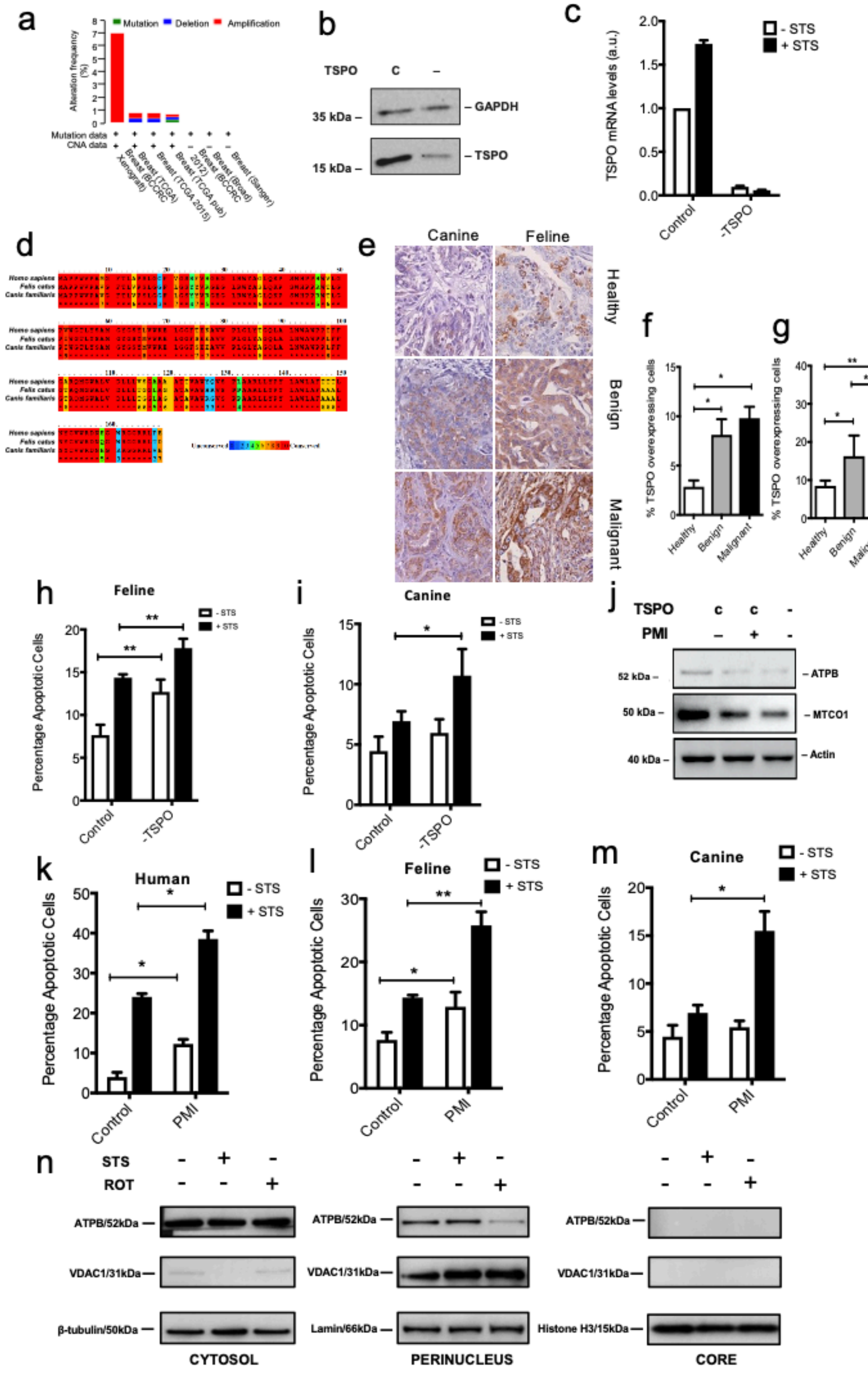

**Supplementary Figure 2**

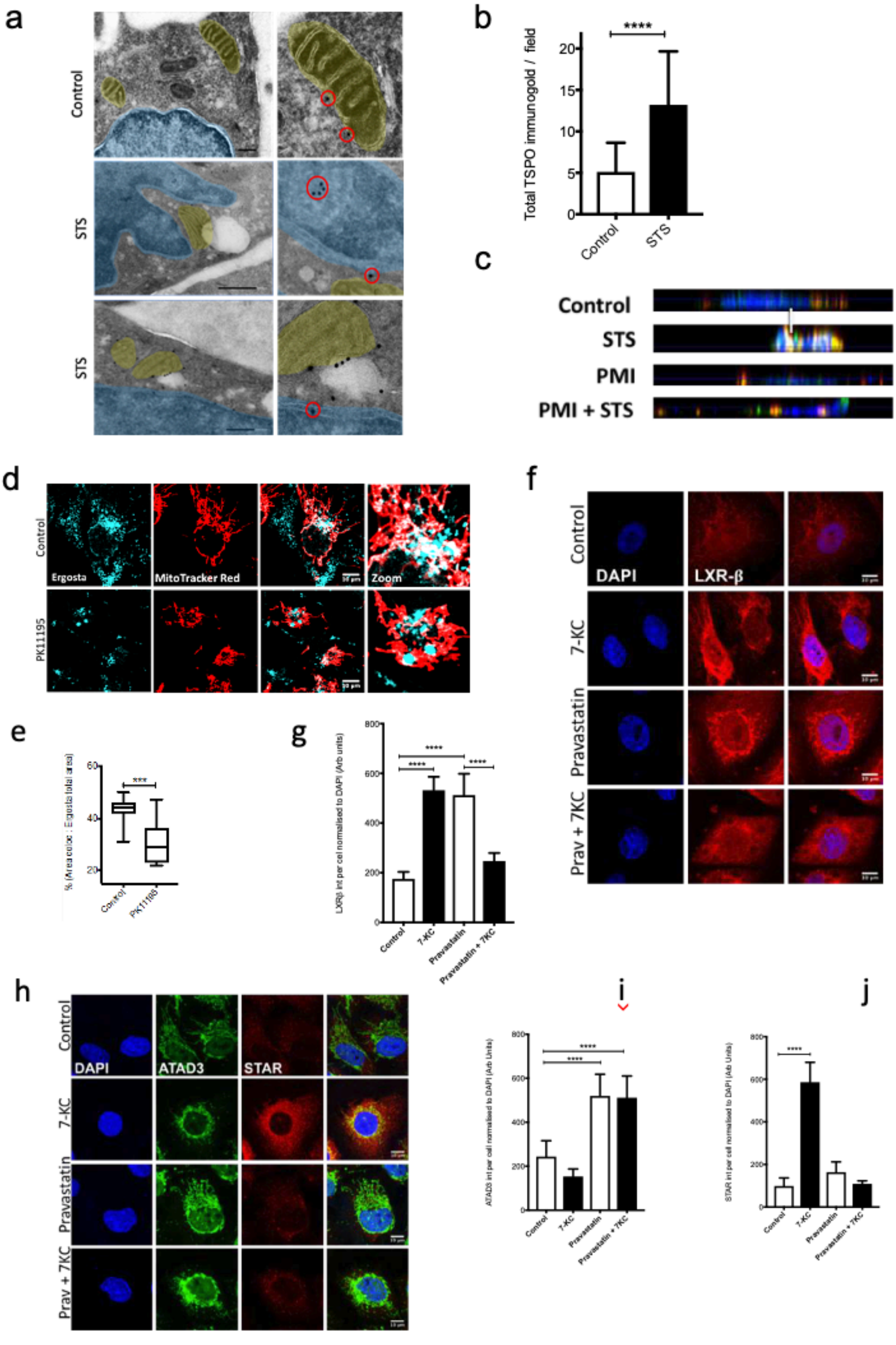

**Supplementary Figure 3**

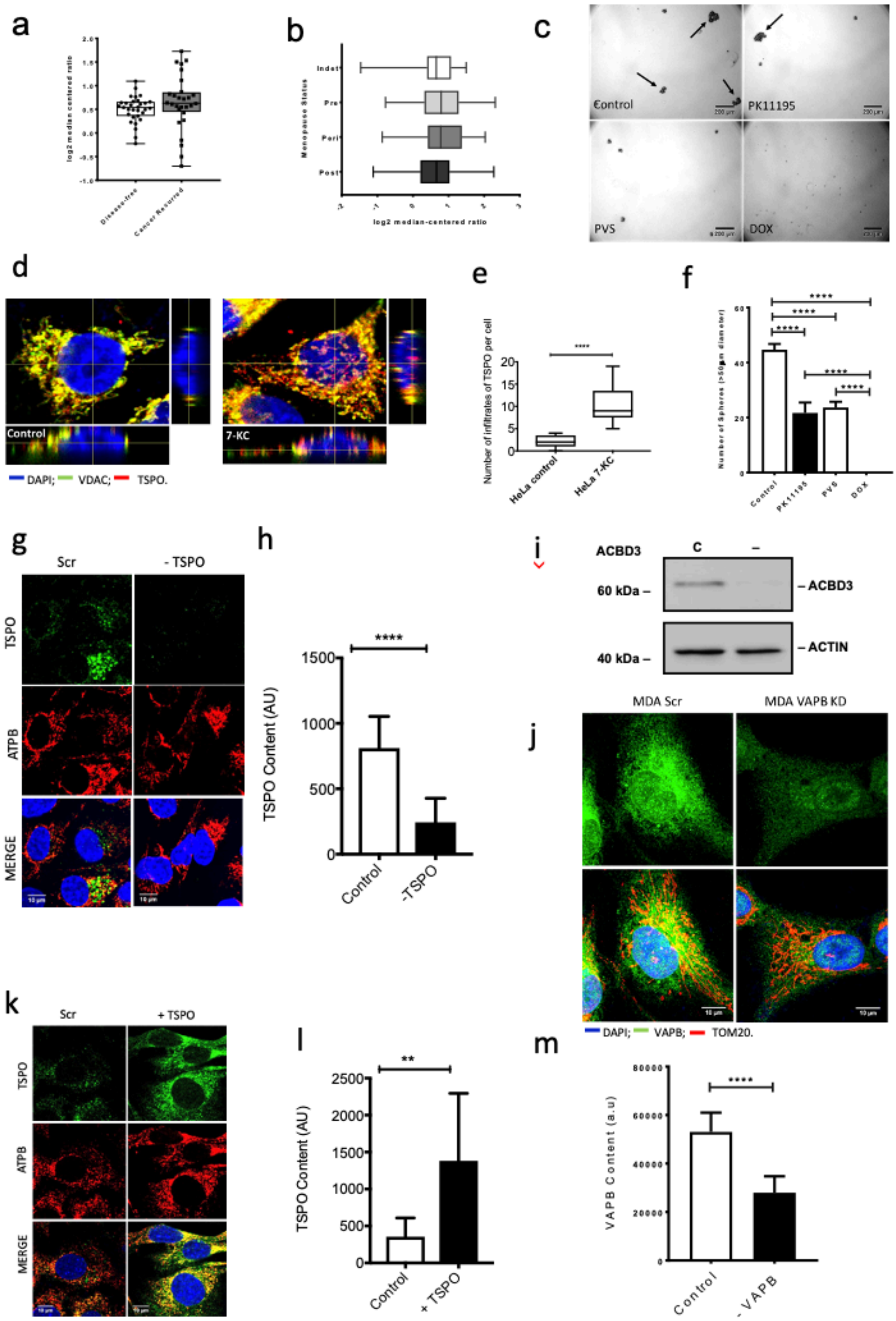

#### Supplementary Figure 4

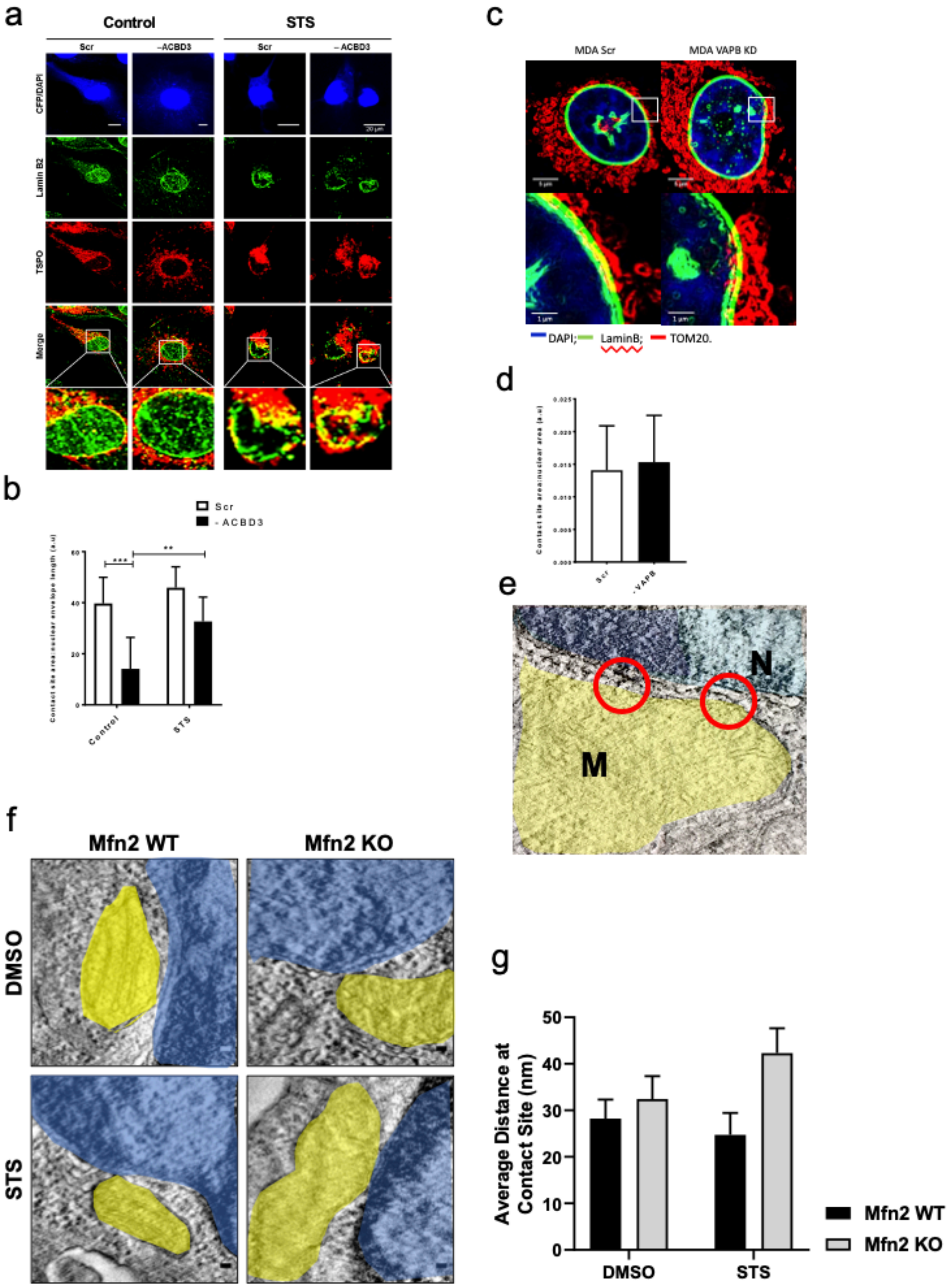

**Legends to Supplementary Figures**

**Supplementary Figure 1: Conservation of TSPO related pathogenesis and resistance across mammalian species.**

(a) Comparison of TSPO degree of mutation in human mammary tumours using microarray data from the TCGA breast cancer database ($p<0.05$).

(b) Western blot showing successful knock down of TSPO in MDA-MB-231 cells using siRNA at 48 hours with the conditions used in experiments.

(c) STS treatment of MDA-MB-231 cells shows increased TSPO mRNA transcript and under siRNA knockdown very low mRNA detected (Control normalised 1; Control +STS 1.174±0.04; -TSPO 0.98±0.01; -TSPO+STS 0.059±0.005; n=3).

(d) Sequence homology shows high level of conservation of the TSPO gene across human, feline and canine genomes.

(e) DAB immunohistochemistry in canine and feline lesions at various stages of disease progression.

(f) Analysis of the DAB immunohistochemistry reveals higher expression in canine mammary cancer tissue compared with healthy, percent TSPO-positive cells significantly more in Canine tissue samples (Healthy 2.85±0.64; Benign 8.12±1.57; Malignant 9.81±1.14; n=20; $p<0.05$).

(g) Analysis of the DAB immunohistochemistry reveals higher expression in feline mammary cancer tissue compared with healthy, percent TSPO-positive cells significantly more in Feline samples (Healthy 8.5±1.4; Benign 16.28±5.4; Malignant2 5.16±3.78; n=4; $p<0.05$, 0.01).

(h) TUNEL assay reports the degree of cell death in feline mammary cancer cells (K48) after STS treatment which highlights how -TSPO cells are more vulnerable to the treatment (Control 7.65±1.2; Control +STS 14.36±0.4; -TSPO 12.69±1.44; -TSPO + STS 17.81±1.09; n=3; $p<0.01$).

(i) TUNEL assay reports cell death in canine mammary cancer cells (CF-41) after STS treatment which highlights how -TSPO cells are more vulnerable to the treatment (Control 4.45±1.19; Control +STS 6.9±0.8; -TSPO 5.9±1.13; -TSPO + STS 10.70±2.2; n=3; $p<0.05$).

(j) Western blot showing the reduction of mitochondrial mass as a result of TSPO reduced expression and PMI treatment as measured by reduced accumulation of mitochondrial proteins ATPB and MTCO1.

(k) Mitophagy inducer PMI exacerbates the impact of STS in human cancer cells and induces significantly higher cell death as quantified by TUNEL assay (Control 3.9±1.2; Control + STS 24.06±.81; PMI 12.28±1.13; PMI+STS 38.54±2.02; n=3; $p<0.05$,0.01).

(l) Impact by PMI on STS induced cell death in feline cancer cells measured by TUNEL assay (Control 7.64±1.2; Control + STS 14.3±.39; PMI 12.9±2.31; PMI+STS 25.78±2.13; n=3; $p<0.05$).

(m) Impact by PMI on STS induced cell death in canine cancer cells quantified by TUNEL assay (Control 4.45±1.19; Control + STS 6.9±0.81; PMI 5.44±0.66; PMI+STS 15.50±2.02; n=3; $p<0.01$).

**Supplementary Figure 2. NAM under apoptotic stimuli, oxidative conditions and patterns of genes expression.**

(a) MDA-MB-231 cells labelled with anti-TSPO immunogold imaged with TEM show that in control (Top panel) TSPO is located on the outer mitochondrial (yellow) membrane whereas after 16 h of treatment (middle panel and lower panel), the nuclear envelope (blue) distorts and TSPO is present on the nuclear envelope (bottom panel) and inside the nucleus (middle panel) as highlighted by red circles (n=2).

(b) Graph shows significant increase in total immunogold labelled TSPO in TEM images of STS treated MDA-MB-231 cells when compared to control counterparts (Control 5.11±3.5; STS 13.21±6.45; n=12; $p<0.001$).

(c) Confocal images showing orthogonal sectioning of MDA-MB-231 cells labelled with DAPI (blue), ATPB (green) and TSPO (red) treated with STS, PMI, STS+PMI showing mitochondrial remodelling and consequent mitochondrial infiltrates under STS treatment.

(d) Confocal images of live staining with fluorescent cholesterol analogue ergosta (dehydroergosterol – cyan) in MDA-MB-231 cells (positive to Mitotracker Red) treated with PK11195 which redistributes the dye away of the focal points.

(e) Confocal images showing increased immunolabeling of oxysterol binding nuclear receptor LXRβ (red) in MDA-MB-231 cells with 7-KC (40 μM for 24 h) treatment as well as Pravastatin treatment (5 μM for 24 hours) which is mitigated when both are applied together indicating the regulation of cholesterol uptake and synthesis

(f) Graph shows ergosta uptake into the mitochondria cell is significantly impaired in PK11195 treated cells compared with control as measured by percent area colocalized with MitotrackerRed (Control 42.79±5.74; PK11195±30.71; n=15; $p<0.005$).

(g) Graph indicating significant increase in LXRβ immunolabeling after treatment with 7-KC (Control 175.5±27.9; 7-KC 532.2±54.12; Pravastatin 512.5±85.9; Pravastatin +7KC 247.2± 32.43; n=8; $p<0.001$)

(h) Confocal images showing no increase immunolabeling of transduceome component ATAD3 (green) in MDA-MB-231 cells with 7-KC (40 μM for 24 h) but higher labelling of Steroidogenic protein STAR. While treatment Pravastatin treatment (5 μM for 24 hours) results in opposite effect. When both are applied together ATAD3 labelling is high whilst STAR is mitigated.

(i) Graph showing increased ATAD3 immunolabeling after treatment with Pravastatin but n

(j) ot 7-KC suggesting it is not involved in oxysterol response unlike TSPO (Control 243.8±72.01; 7KC 153.6±33.9; Pravastatin 519±98.19; Pravastatin + 7KC 511.5±98.64; n=8; p<0.001)

(k) Graph showing increased STAR immunolabeling after treatment with 7-KC but not Pravastatin suggesting it is involved in oxysterol response like TSPO, however this is mitigated with pre-treatment of pravastatin (Control 98.5±37.844; 7KC 586.4±93.16; Pravastatin 163±48.471; Pravastatin + 7KC 109±14; n=8; p<0.001)

**Supplementary Figure 3. Pattern of TSPO accumulation in relapsed breast cancer patients and in HeLa cells treated with 7-KC**

(a) Comparison of TSPO expression in humans grouped by estrogen positive breast cancer recurrence following tamoxifen using microarray data from the Ma Breast 2 study (Ma et al., **https://doi.org/10.1016/j.ccr.2004.05.015**). Line at median, error bars show min to max, n=60.

(b) Comparison of TSPO expression grouped by menopausal status in human mammary tumours using microarray data from the TCGA breast cancer database. Indeterminate n=34 (Neither Pre- or Post-Menopausal). Pre n=108 (<6 months since last menstrual period and no prior bilateral ovariectomy and not on oestrogen replacement). Peri n=14 (6-12 months since last menstrual period). Post n=238 (Prior bilateral ovariectomy or >12 months since last menstrual period with no prior hysterectomy). Line at median, error bars show min-max. p<0.01.

(c) Images of mammosphere formation assay showing that both PK11195 and PVS treatment significantly reduce the ability of MDA-MB-231 cells to grow in an anchorage-independent manner. A sphere is defined as a cluster of cells with a diameter greater than 50μm in length. DOX was used as a positive control.

(d) Confocal images showing orthogonal sectioning of HeLa cells labelled with DAPI (blue), ATPB (green) and TSPO (red) treated with 7-KC showing upregulation of TSPO and consequent mitochondrial infiltrates under 7-KC treatment.

(e) Calculation of the mitochondrial infiltrates into the nucleus measured in orthogonal images of 7-KC treated HeLa cells (Control 2.222±0.4648 n=9; 7-KC 10.44±1.454 n=9).

(f) Quantification of number of spheres (>50μM diameter) per well following equal seeding. Results reveal a role of TSPO and cholesterol in breast cancer invasiveness and progression (Control 46.752±42.588; PK11195 25.456±17.884; PVS 25.752±21.588; n=3 $p<0.001$).

(g) Immunofluorescence images showing successful knockdown of TSPO in MDA-MB-231 cells using siRNA at 48 hours prior to fixing.

(h) Quantification of TSPO content in immunofluorescence images, TSPO levels are significantly lower in –TSPO MDA-MB-231 cells (Control 1052.7±30.9; -TSPO 427.3±60.3; n=10 $p<0.001$)

(i) Western blot showing successful knock down of ACBD3 in MDA-MB-231 cells using siRNA at 48 hours with the conditions used in experiments.

(j) Immunofluorescence images showing successful knockdown of VAPB in MDA-MB-231 cells using siRNA at 48 hours prior to fixing.

(k) Immunofluorescence images showing successful overexpression of wildtype human TSPO in mouse embryonic fibroblasts (MEF) following plasmid transfection.

(l) Quantification of TSPO content in immunofluorescence images, TSPO levels are significantly higher in +TSPO MEF cells (Control 608.1±101.5; +TSPO 2295.33±468.7; n=10 $p<0.01$).

(m) Quantification of VAPB content in immunofluorescence images, TSPO levels are significantly lower in –VAPB MDA-MB-231 cells (Control 61004±45088; - VAPB 34741±20541; n=10 $p<0.001$).

**Supplementary Figure 4. The formation of NAM is independent from MAM**

(a) Confocal images of wild-type and ACBD3 knock-down MDA cells. Cells were co-transfected with cyan fluorescent protein (CFP, blue) and immunostained for lamin B2 (green) and TSPO (red) following treatment with DMSO-vehicle or STS.

(b) Quantification of lamin B2 and TSPO co-localisation from confocal images. Knocking-down ACBD3 in control cells reduces mitochondria-nucleus interactions in untreated cells. However, following STS treatment, loss of ACBD3 does not significantly reduce the co-localization between TSPO and lamin B2 in comparison to wild-type cells.

(c) Confocal images of DAPI (blue), Lamin B (Green) and mitochondrial TOM20 (red). Silencing known tether protein VAPB in MDA-MB-231 cells does not prevent the formation of contact sites observed by co-localisation (yellow) of the nuclear envelope and mitochondria.

(d) Quantification of lamin B and TOM20 co-localisation normalised to nuclear area in scrambled siRNA and VAPB knockdown MDA-MB-231 cells. (Control 0.202081±$7.259 \times 10^{-3}$; -VAPB 0.022475±$8.125 \times 10^{-3}$; n=15, ns)

(e) Representative TEM micrograph of a mitochondria-nucleus contact site in MEFs,

Indicating the presence of a specialized tethering structure between the mitochondrial and nuclear membranes.

(f) Representative TEM micrograph of a mitochondria-nucleus contact site in Mfn2 ko MEFs control or STS treated.

(g) Quantification of minimum distance between mitochondria and the nucleus in control and STS treated Mnf2ko MEFs (n=3).
